# CRAM 3.1: Advances in the CRAM File Format

**DOI:** 10.1101/2021.09.15.460485

**Authors:** James K Bonfield

## Abstract

**Motivation:** CRAM has established itself as a high compression alternative to the BAM file format for DNA sequencing data. We describe updates to further improve this on modern sequencing instruments.

**Results:** With Illumina data CRAM 3.1 is 7 to 15% smaller than the equivalent CRAM 3.0 file, and 50 to 70% smaller than the corresponding BAM file. Long-read technology shows more modest compression due to the presence of high-entropy signals.

**Availability:** The CRAM 3.0 specification is freely available from https://samtools.github.io/hts-specs/CRAMv3.pdf. The CRAM 3.1 improvements are available from https://github.com/samtools/hts-specs/pull/433, with OpenSource implementations in HTSlib and HTScodecs.

**Supplementary information:** Supplementary data are available online

## 1 Introduction

It has been well established that the growth in genomic sequencing data is challenging (Stephens *et al*., 2015). The earlier file formats of SAM and BAM (Li *et al*., 2009) were appropriate for the era, but better techniques were soon required. The notion of reference-based compression, storing only the differences between DNA sequence fragments and the reference they have been aligned against, was proposed (Fritz *et al*., 2011). Fritz *et al*. also proposed techniques for efficient encoding of unaligned data by the use of sequence assembly to generate consensus sequences, which may then be used as the reference sequence to compare against. This work lead to the development of CRAM by the European Bioinformatics Institute (Cochrane *et al*., 2013).

The primary goals of CRAM were a reduction in storage requirements, while maintaining direct compatibility with BAM, permitting lossless round trips. All data representable in BAM is also available in CRAM. This includes the SAM header, which is the same format in CRAM, and the optional auxiliary key-value “tags”. These annotations are defined by a shared SAMtags specification (samtools.github.io/hts-specs/SAMtags.pdf).

Although reference compression is where the original work focused, it is wrong to assume that this is the primary reason for CRAM’s reduced file size. BAM serialises all data together (first name, chromosome, position, sequence, quality and auxiliary fields, then second name, chromosome and so on). This leads to poor compression ratios as names, sequences and quality values all have very different characteristics. CRAM has a column-oriented approach where a block of names are compressed together, or a block of qualities. Each block can be compressed with an algorithm specific to that data type. This leads to significantly reduced file sizes and is often the biggest factor in file reduction.

The first tool implementing CRAM (then version 1.0) was CRAMtools (Vadim Zalunin, 2011, unpublished), written in Java. The Scramble tool (Bonfield, 2014) was the first C implementation and lead to a specification tidy-up producing CRAM 2.0 in 2013. HTSlib (Bonfield et al., 2021) gained CRAM support shortly after. CRAM 3.0 appeared a year later in 2014, with some additional compression codecs including the rANS entropy encoder (Duda, 2013) and LZMA (Lempel Ziv Markov-chain Algorithm, Igor Pavlov, 1998, unpublished). More implementations of CRAM have since appeared, written in JavaScript (Buels et al., 2019) and Rust (https://github.com/zaeleus/noodles). Many more programming languages support CRAM via bindings to one of these existing implementations.

The CRAM specification is now maintained by the Global Alliance for Genomics and Health (GA4GH: https://www.ga4gh.org/cram/). It ties in with a number of other GA4GH standards and protocols which further extend the features and capabilities. Reference sequences may be obtained either via local files or using a refget server (Yates et al., 2021). CRAM files can be streamed remotely using the htsget protocol (Kelleher et al., 2019), and they may be encrypted using Crypt4GH (Senf et al., 2021).

Since 2014 CRAM has been very stable, but a lot has changed data-wise. Illumina’s quality values have been successively quantised from 40 discrete values, to 8, and now with NovaSeq to 4 (Illumina, 2012). We have also seen the rise of long-read technologies and more complex auxiliary data types being embedded in the files. As the data changes, so too should the encoding and compression methods available to the format. Methods such as Run Length Encoding (RLE) were considered and explicitly rejected as unhelpful in the original CRAM development, but now these same techniques can be beneficial. CRAM 3.1 is the first major update to CRAM since 2014. It keeps the underlying format unchanged, but adds new compression codecs.

With large data volumes comes large processing requirements. By default CRAM optimises for a balance between CPU cost, file size and granularity of random access. However the option of higher memory and CPU requirements for long-term archival is still worthy of consideration so CRAM 3.1 also improves support for archival modes.

At the time of writing CRAM 3.1 is in draft. Implementations of the new codecs exist in C (HTSlib, SAMtools and Scramble) with a JavaScript proof of concept.

## 2 Methods

The basic structure of CRAM can be seen in Figure 1. It starts with a header matching the SAM specification, although it mandates the use of MD5sums on reference sequence lines for data provenance and to ensure correct decoding.

**Fig. 1.**
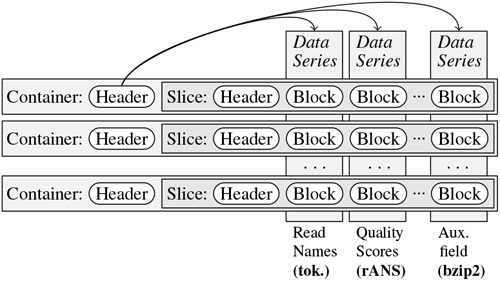
Logical layout of a CRAM file, showing containers and slices as rows, and data-series as columns. Random access is possible on rows, with rapid filtering (discarding) of columns.

CRAM’s records are broken down into data series, loosely fitting the columns in a SAM file, such as alignment position, quality values or CIGAR string components. Each auxiliary tag also gets its own data series.

The CRAM header is then followed by a series of containers, which in turn hold slices and data blocks within them. The container header consists of meta-data describing how and where each data series is encoded. The slices are collections of alignment records, applying the encoding rules described by the container to the records and storing the result in the requested blocks. The blocks are then compressed using their own selected compression algorithms. It is these algorithms which have been added to in CRAM 3.1. Slices may be of any size, but the HTSlib implementation defaults to 10,000 records, or fewer if long reads are present.

This offers great flexibility to the encoder, meaning that over time encoder improvements may yield smaller files. For example the first 10 million reads of a NovaSeq alignment in CRAM 3.0 format produced by HTSlib 1.2 takes up 199.6MB. The latest HTSlib 1.13 encodes the same file in 195.2MB and Picard 2.25.7 with default options uses 254.1MB. All of these files are compatible and have the same choice of codecs available.

As it is possible to store each data series in its own block, this permits selective decoding where only specific types of data need to be decoded. This can be of great benefit to certain algorithms. For example the “samtools flagstat” command gives a summary of SAM FLAG bit frequencies (Danecek *et al*., 2021). Although it does not need to know about sequences, quality values or read identifiers, with a BAM file it is still required to decompress this data due to the serial nature of the format. With CRAM it only decompresses the data series required. Consequentially “samtools flagstat” on the NA12878 Platinum Genomes file takes 7m35s on the CRAM file and 22m52s on the BAM file (with neither file in disk cache).

The NM and MD SAM auxiliary tags have special handling within CRAM. As they describe the difference between an aligned sequence and the reference and we are typically doing reference based compression, they may be omitted and generated on-the-fly during decode. If these values are found to be in error then they can either be corrected, or if we wish to have bug-compatible data then the (incorrect) values may be stored verbatim in the CRAM file.

Each slice can optionally also contain a copy of the reference used for that genomic region. This permits CRAM to do reference-based compression while removing the dependency on external data files. For deeply covered regions this does not have a significant impact on compression ratios. This embedded reference could be a consensus rather than the official external reference, offering the potential for improved compression via fewer sequence differences. However doing so means storing NM and MD verbatim in regions where consensus and reference differ, negating most of the gains. Note this problem is resolved in Deez (Hach *et al*., 2014) by using a two-level delta (sequence to consensus and consensus to reference), and may be considered for a future CRAM update.

The compression codecs permitted in CRAM 3.0 are three external general compression tools-deflate (Deutsch and Gailly, 1996), bzip2 and LZMA – and the rANS entropy encoder. The entropy encoder uses static frequencies, written at the start of the block, which can be either Order-0 or Order-1 metrics. An example of Order-0 frequencies is the observation that the letter “u” accounts for 3% of the total letter usage in English, while an Order-1 observation is that “u” occurs nearly 100% of the time following the letter “q”.

The current official CRAM format is 3.0, but CRAM 3.1 has been a draft standard since 2019. The layout of the CRAM format is unchanged, but new custom compression codecs have been added. These include:

### Improved rANS with data transformations

The reduction in the count of discrete quality values in Illumina data from 40 (HiSeq) to 4 (NovaSeq) has meant that some simple data transformations can reduce both data size and time to encode. The newer rANS codec has bitpacking, for example mapping 4 distinct quality values to 2 bits each and storing 4 per byte, and run-length encoding. Additionally the 4 rANS states used in CRAM 3.0 can now be expanded to 32 states permitting the use of SIMD instructions for accelerated encoding and decoding.

### Adaptive arithmetic coder

This is a byte wise arithmetic coder with adaptively updated frequencies. This helps for data types with non-stationary probability distributions, but it has a higher CPU overhead. It also includes the data transformations used in the updated rANS codec. This coder is internally used by the FQZComp quality coder and optionally by the name tokeniser. It has order-0 and order-1 models, but it could trivially be extended to support higher order models if deemed necessary in the future.

### FQZComp quality encoder

This is a generalised version of the quality model used in the FQZComp tool (Bonfield and Mahoney, 2013). It is limited to a maximum of 16 bits of context in order to permit rapid model tuning and to make it appropriate for a more random-access oriented file format. The context is applied to an adaptive arithmetic encoder. This makes it slower than rANS and more suitable to data archival. The construction of this context is flexible and described within the file format, offering great opportunity for learning and tuning to a specific data set. Data available for model context generation include the previous quality values, the position along the current read, a cumulative delta of successive quality values since the start of each read, bits indicating reverse complement or duplicate, and a general context selector copying from bits stored in the file (which may be used to split by average quality value or x/y location extracted from the read name for example). Values may also be transformed through lookup tables prior to context generation. An example FQZComp configuration is shown in Figure 2.

**Fig. 2.**
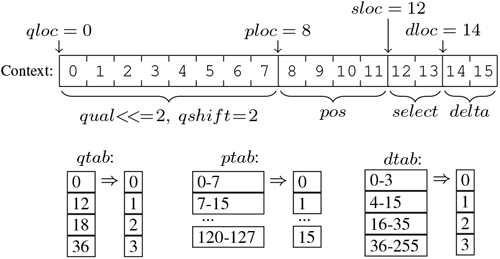
An example FQZComp configuration describing how previous quality values, the position in the sequence, a running sum of the delta between qualities and a generic model selector can be combined with lookup tables to generate a context model.

### Read name tokeniser

The read identifiers are often highly structured, such as “HSQ1004:134:C0D8DACXX:4:2107:20375:180666”. Much like how CRAM separates the primary SAM fields into columns, the name tokeniser separates the components of a read names and compares against a component in a previous name. Each component column is then serialised with optional string or numeric deltas and compressed using either the static rANS or adaptive arithmetic encoders.

CRAM also permits some controlled data loss. Read names may be discarded, with new names generated during decode. Read pairing within a slice is encoded by explicit links between records, so the generated names are still in pairs. Quality values may also be omitted, with only specific sites being stored. However this is not the recommended approach to quality value reduction. It is possible to reduce the entropy of quality value either by smoothing methods such as P-block (Cánovas et al., 2014) or sitespecific quality reassignment via CALQ (Voges et al., 2018) or Crumble (Bonfield *et al*., 2019). These modified quality strings are then much more compressible, particularly, when combined with the newer RLE rANS methods. As such we view quality loss best dealt with as a topic external to the file format.

The primary focus of CRAM is with sorted aligned data. However SAM, BAM and CRAM all support unaligned data too and the addition of both per-file and per-read meta-data arguably make these a superior format to using FASTQ. There are many FASTQ compression tools which offer superior ratios to unaligned CRAM, but our approach to FASTQ is primarily as a transitional format between sequencing and either alignment or assembly rather than as a suitable long-term archival format. That said, combining an approximate rapid sequence aligner with CRAM can be used to reduced data size. Examples of this using SNAP (Zaharia *et al*., 2011) are in the Supplementary Material.

CRAM also permits aligned non-position-sorted data, such as by read name order. This may use either reference-based or reference-less encoding, with the latter sometimes being a more time and memory performant option when dealing with very large genomes.

## 3 Results

This manuscript is on the CRAM format rather than a specific implementation, however it is not possible to analyse the performance of the format without evaluating an implementation. We use SAMtools and HTSlib 1.13. Where possible, we also compare against Deez, MPEGG and Genozip (Lan et al., 2021). The GenomSys MPEG-G tool is not freely available so figures reported are taken from their paper and its references. Note their CRAM figures significantly differ to ours. We requested clarification from the authors, but some of these differences remain unexplained.

Benchmarks for the new codecs in isolation, outside of CRAM, can be seen in the tables below.

Table 1 shows entropy encoder speeds for the first 1 million records from NovaSeq data. Speeds on HiSeq 2000 qualities are listed in the Supplementary Material. Also shown are other static frequency entropy encoders including Zlib’s Huffman encoder and the Finite State Entropy (FSE) implementation used in Zstd (Collet, 2021). These only support Order-0 encoding and Huffman is unable to encode the skewed 4-quality NovaSeq data efficiently.

**Table 1.** NovaSeq quality: entropy encoder speeds.

| Program | Option | Size(MB) | Enc(MB/s) | Dec(MB/s) |
| --- | --- | --- | --- | --- |
| Zlib | Huffman | 21.48 | 151.3 | 365.4 |
| FSE | tANS | 9.90 | 435.2 | 531.1 |
| FSE | Huffman | 20.74 | 709.8 | 1302.9 |
| rANS4x8 | O0 | 9.91 | 391.4 | 531.1 |
| rANS4x16 | O0 | 9.90 | 446.5 | 823.7 |
| rANS32x16-AVX2 | O0 | 9.92 | 617.5 | 1609.3 |
| rANS4x8 | O1 | 9.14 | 271.2 | 406.8 |
| rANS4x16 | O1 | 9.14 | 285.4 | 551.7 |
| rANS32x16-AVX2 | O1 | 9.16 | 356.7 | 1091.7 |
| rANS32x16-AVX512 | O0,PACK+RLE | 9.18 | 583.6 | 1378.3 |
| rANS32x16-AVX512 | O1,PACK+RLE | 8.26 | 489.3 | 1123.1 |
| arith | O0 | 9.83 | 120.0 | 94.5 |
| arith | O0,PACK+RLE | 9.16 | 220.0 | 188.3 |
| arith | O1 | 9.12 | 105.3 | 91.3 |
| arith | O1,PACK+RLE | 8.12 | 156.5 | 115.1 |
| GABAC-app | -d15 -s1 | 8.40 | 5.0 | 8.0 |

Block size 1MB, except FSE Huffman which used 128KB.

This table also shows performance of the adaptive coder along with the Genomic Adaptive Binary Arithmetic Coder (GABAC) (Voges et al., 2020) used in the OpenSource Genie (Bliss *et al*., 2018) reference implementation of MPEG-G (Voges *et al*., 2021). This latter provides a reasonable ratio for the low-entropy NovaSeq data set, but is two orders of magnitude slower than the faster methods. Note this may not be indicative of a well optimised GABAC implementation.

Table 2 shows the performance of two predefined FQZComp configurations on high-entropy HiSeq 2000 quality values and low-entropy NovaSeq quality values, compared against libbsc. It can be seen that the choice of model configuration can be critical. This particular set of HiSeq 2000 data has some highly erroneous cycles, so using more bits to track position within the read is very productive. Additionally this model utilises the embedded selector bits to separate data by READ1 and READ2 flags, slightly improving the compression of the NovaSeq data too.

**Table 2.** Quality value FQZComp performance.

| Program | Option | Size(MB) | Enc(MB/s) | Dec(MB/s) |
| --- | --- | --- | --- | --- |
| *NovaSeq qualities* |  |  |  |  |
| bsc | -m0e2tTp | 7.72 | 19.40 | 35.48 |
| FQZComp | -s0 | 7.27 | 28.57 | 51.80 |
| FQZComp | -s1+read 1/2 | 7.21 | 21.43 | 26.74 |
| *HiSeq 2000 qualities* |  |  |  |  |
| bsc | -m0e2tTp | 43.8 | 6.75 | 9.11 |
| FQZComp | -s0 | 42.5 | 15.84 | 17.82 |
| FQZComp | -s1+read 1/2 | 31.3 | 13.29 | 14.91 |

Compression of NovaSeq and HiSeq 2000 quality values using libbisc and CRAM 3.1's FQZComp, in 10 blocks of 100,000 records.

Table 3 shows the performance of the name tokeniser on 10 blocks each containing 100,000 NovaSeq read names. The tokeniser is considerably smaller than general purpose tools on the more predictable name-sorted data. With chromosome and position sorted data, which scrambles the name ordering, the tokeniser is only beaten by the much slower “mcm” tool.

**Table 3.** NovaSeq read name compression.

| Program | Name sorted |  |  | Position sorted |  |  |
| --- | --- | --- | --- | --- | --- | --- |
|  | Size | Enc(MB/s) | Dec(MB/s) | Size | Enc(MB/s) | Dec(MB/s) |
| bzip2 | 2.52 | 32.08 | 222.11 | 6.18 | 15.97 | 137.88 |
| gzip -12 | 2.14 | 3.78 | 250.59 | 5.50 | 2.80 | 241.93 |
| bsc -m5e1tT | 1.68 | 27.61 | 23.07 | 3.99 | 22.59 | 19.53 |
| xz -9 | 1.31 | 2.50 | 74.98 | 4.87 | 1.70 | 65.31 |
| mcm -m7 | 1.22 | 3.07 | 3.21 | 3.43 | 2.69 | 2.78 |
| tok3 -7 | 0.89 | 15.46 | 48.54 | 3.55 | 10.00 | 69.44 |
| tok3 -19 | 0.88 | 8.82 | 41.59 | 3.48 | 4.84 | 39.13 |

Performance of the read name tokeniser on name sorted and chromosome / position sorted NovaSeq reads. Size is shown in MB.

Overall CRAM performance is not a single metric as it permits user-adjustable trade-offs between speed, size and granularity of random access, multiple points have been plotted for each CRAM version. The CRAM benchmarks are from the released version of SAMtools 1.13. Note this does not yet include the SIMD-vectorised rANS entropy encoder and is using the 4-way Scalar implementation. Results using a vectorised build of SAMtools are presented in the Supplementary Material.

All decode timings are a read and complete decode of the files, with data discarded where possible. Encode timings are conversion from compressed BAM to a new file format. Full benchmarks and details are available in the supplementary material. The encoder and decoder were given 12 threads, running on an Intel Xeon CPU E5-2660 running at 2.20GHz. The quoted MPEG-G timings used 12 threads on an Intel Xeon E5-2670 at 2.6GHz.

Figure 3 shows the results for the Illumina HiSeq 2000 (ERR194147), Illumina NovaSeq (ERR3239334) and PacBio CLR (ftp://ftp.1000genomes.ebi.ac.uk/vol1/ftp/technical/working/20131209_na12878_pacbio/si/NA12878.pacbio.bwa-sw.20140202.bam). All three are Whole Genome Shotgun (WGS) libraries of NA12878.

**Fig. 3.**
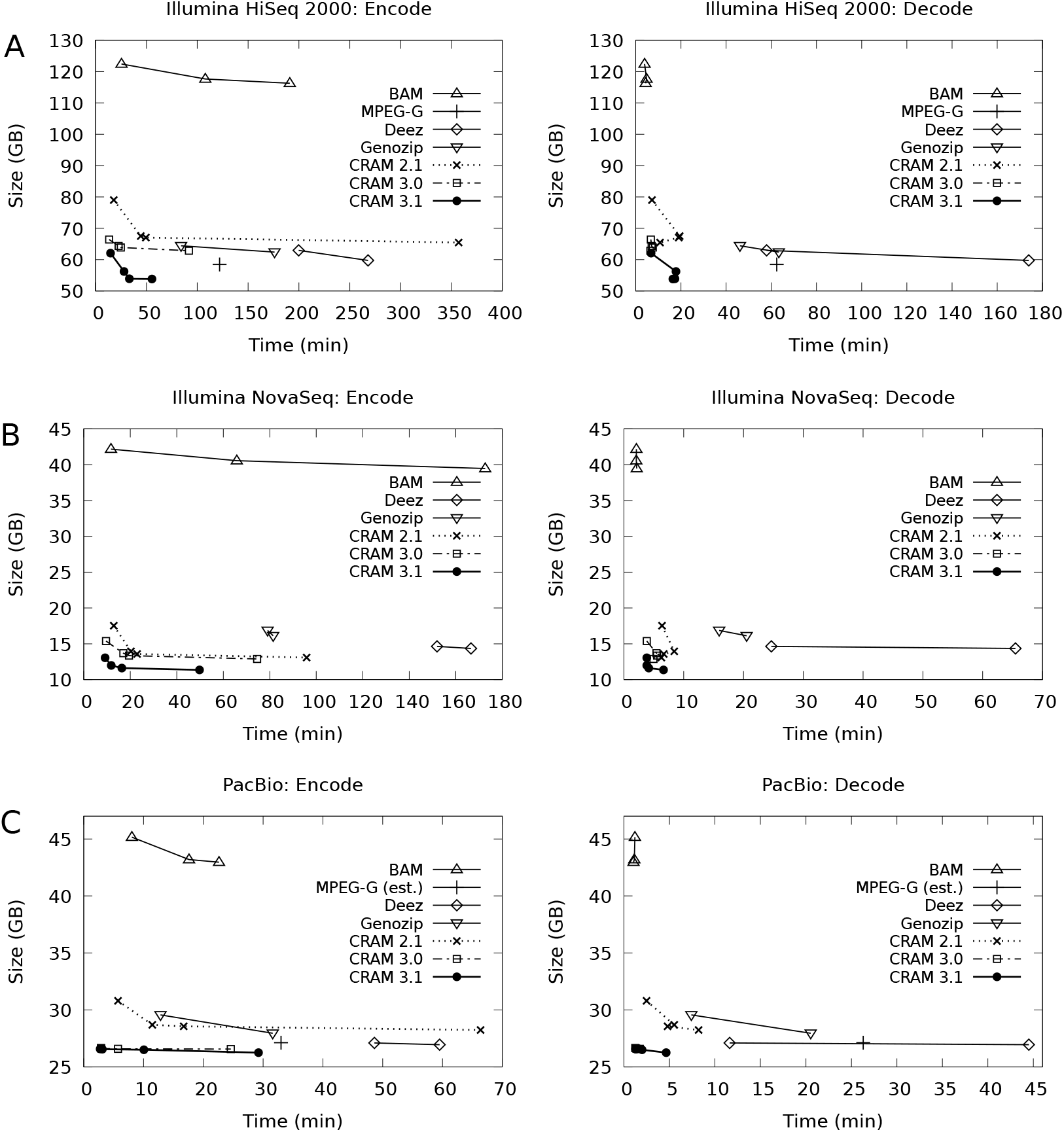
Benchmarks of aligned data formats using 12 threads. MPEG-G figures are taken from the Voges et.al paper, with “MPEG-G (est.)” possibly using a slightly different input file. (See text.)

Figure 3 part A shows the encode and decode speeds against file size for the Illumina HiSeq 2000 data (ERR194147). This is also MPEG-G data-set 02. Some formats have multiple points, linked together by a line. This represents different compression profiles used by the tools. In BAM this adjusts the libdeflate compression levels to 6, 10 and the maximum of 12.

CRAM charts show profiles “normal”, “small”, “archive” and “archive” at compression level 9. The default “normal” profile only uses the rANS codec and Deflate. The “small” profile increases the block size and enables Bzip2 compression and in CRAM 3.1 the FQZComp codec. The “archive” profile further increases block size, boosts the deflate compression level and in CRAM 3.1 enables the adaptive arithmetic coder. At compression level 8 and above the LZMA codec is also enabled for all CRAM versions. This has a significant cost to compression time so is often not a good speed / size trade-off, but is still fast at decoding.

It can be seen that CRAM 3.1 offers a similar leap over CRAM 3.0 compression ratios that it did in turn over CRAM 2.1. CRAM 3.1 is between 7 and 16% smaller than CRAM 3.0 at the equivalent profile, while being similar on speed except for “small” and “archive” decoding times. All CRAMs except those using LZMA encode faster than their BAM equivalents, while saving up to 56% storage, although BAM is quicker to decode. This is compatible with our design goal of being similar speed to BAM.

It is unclear whether the published MPEG-G benchmarks include auxiliary tags, but these only account for under 1% of this file. CRAM 3.1’s compression profiles straddle the MPEG-G file size, with the “small” CRAM profile being both smaller and 4 times faster than MPEG-G. However note the MPEG-G benchmarks were not performed by us, so there may be differences in measuring techniques.

Deez compression ratios are between CRAM 3.0 and CRAM 3.1 and a little behind MPEG-G. It is the slowest tool, although it should be noted that despite being given 12 threads it typically only utilised 2. Total CPU usage was less than MPEG-G. Genozip was also much slower than CRAM 3.1 while being larger, similar in size to CRAM 3.0 archive mode.

Figure 3 part B shows the same NA12878 sample sequenced using an Illumina NovaSeq instrument (ERR3239334). The improvements of CRAM over BAM here are more marked, due to the higher compressibility of the quantised quality values. The gains from CRAM 3.0 to CRAM 3.1 are also greater, with the “normal” profile being 16% smaller. The CRAM 3.1 files are between 3.1 and 3.5 times smaller than BAM.

As before, Deez is slow and with this file does not match CRAM 3.0 for file size. No MPEG-G results are available for this data set, but the published results for the NovaSeq MPEG-G data set 37 implies good compression performance with a size similar to CRAM 3.1 archive mode. Genozip performed poorly, with file sizes larger than CRAM and considerably slower.

Figure 3 part C shows an aligned PacBio CLR file, also for NA12878. This is the MPEG-G data set 03. Published results are available for MPEG-G on this data, but we were only able to get our CRAM files to match their earlier published results (MPEG document M56361) by removing secondary alignments and discarding auxiliary tags. Hence for comparison purposes we applied these transformations to the downloaded BAM before performing this benchmark. We have been unable to verify with the authors if this is the correct procedure used in the MPEG-G publication, so it is listed here as “MPEG-G (est.)”.

This data set shows a very minimal change between CRAM versions and compression profiles. The file is dominated by the quality values, which are largely uncompressible due to having a big range of discrete values (0 to 93) with very little correlation between successive values. Nevertheless it is evident that CRAM saves a significant portion over BAM and CRAM 3.0/3.1 is a big improvement on the historic CRAM 2.1. The assumed MPEG-G size, Deez and Genozip are all larger than CRAM 3.0 while being significantly slower.

## 4 Discussion

CRAM has achieved the goals of providing a space-efficient alternative to BAM, while not suffering a significant time penalty. CRAM files in the European Nucleotide Archive significantly outnumber BAMs (personal communication) and it has seen wide adoption at many other sites. While some other tools are now smaller than CRAM 3.0 on some data files, this typically comes at a heavy cost in CPU. We demonstrate with CRAM 3.1 that changing just one component of the CRAM format, the compression codecs available, is sufficient for CRAM to be competitive on size once more while not sacrificing its speed advantage. We also expect CRAM 3.1 to continue to improve as we learn how best to tune each codec, in particular the selection of the optimal FQZComp models.

However there are wider format adjustments that could be made over and above adding new compression codecs, which will be addressed in CRAM 4.0. While an early draft of this already exists, it is likely to undergo further revisions. Improvements include migration of the rANS PACK and RLE filters to the CRAM slice encoding methods with additional transformations such as integer delta encoding (useful for Oxford Nanopore Technology signal data) and an LZP step to help reduce repetitive auxiliary tags and to remove duplication in quality values due to secondary alignments. Potentially further custom codecs may also need to be developed for the most expensive auxiliary tags, such as breaking down comma-separated tag formats (e.g. SA:Z) into components similar to the read name tokeniser. The general purpose compression interfaces used may also be reconsidered, such as replacing Deflate with Zstd and Bzip2 with Libbsc (Grebnov, 2011). There is scope for moving some compression meta-data, such as the order-1 frequency tables in rANS, out of the data block produced by compression codecs and into the container header. This could permit finer grain random access with many slices per container. Finally improvements can be made to how embedded consensus sequences are handled, perhaps using a two-step delta as implemented in Deez.

It is evident however that some data does not gain compression with CRAM 3.1 due to the inherent randomness of the data held within, and no format improvement is likely to solve that. This is particularly true for some long-read technologies. We question the requirement to have so many distinct quality values in PacBio and ONT data and suggest lossy compression may be suitable for these data sets. We feel that this is best researched and addressed by the sequencing manufacturers and urge them to consider ways to reduce the data footprint of their output files.

An implementation of the CRAM 3.1 and 4.0 draft standards may be found in HTSlib (https://github.com/samtools/htslib) and HTScodecs (https://github.com/samtools/htscodecs).

## Acknowledgements

Vadim Zalunin (European Bioinformatics Institute) was instrumental in producing the first production-ready implementation of CRAM, upon which the rest of this work sits. We would also like to thank Chris Norman (Broad Institute) for his work as a co-maintainer of the CRAM specification and continued support for CRAM within htsjdk. The Order-0 rANS codec was based on work by Fabian Giesen and the arithmetic coder includes work by Eugene Shelwien. Andrew Whitwham proof read the paper and Rob Davies performed code reviews, both from Wellcome Sanger Institute.

