## Supplementary Material for "CRAM 3.1: Advances in the CRAM File Format"

### CRAM 3.1: Advances in the CRAM File Format Supplementary Material

James Bonfield

October 11, 2021

#### Testing methodology

Encoding times were performed using SAMtools, Deez and Genozip converting from compressed BAM to the desired format. The decoding time of BAM is one of the fastest and is typically much faster than encoding time, so the impact of this on the results is low. Additionally we felt that compressed BAM offered the best format to read from as it reduced potential I/O bottlenecks, particularly when multi-threading.

Decode times for SAMtools were performed using "samtools view -f 0xffff". This reads the data and filters out anything that doesn't have every bit-FLAG set, which is all reads. Hence there is no output and the time is dominated by decode only. Note for comparison against other tools, testing the fastest modes of CRAM decoding shows converting to uncompressed BAM adds 25% additional CPU cost over simply discarding the data. This further reduces in percentage with the slower CRAM profiles.

While we attempted the same filtering with Deez, it cleverly detected it could omit most of the work so the times were almost instantaneous. Therefore Deez decode timings unfortunately include outputting the data to stdout, which was then redirected to /dev/null to avoid file I/O costs. Similarly for Genozip we redirected output to /dev/null.

We do not know how the MPEG-G timings were performed other than the system they used (Intel Xeon E5-2670 at 2.6GHz).

Programs were given 12 threads for the aligned data and 8 threads for unaligned. The reason for difference is due to the availability of the MPEG-G benchmarks to compare against and we attempted to use a like for like test.

Times were measured using either "perf stat" or the unix "time" command, to record separately the user+system CPU time and Real (elapsed) time. The main paper shows the real time, but we also include CPU times below.

Unless specified otherwise, the machine used for benchmarking was an 16-core Intel Xeon CPU E5-2660 running at 2.20GHz with 2 hyper-threads on each core. The operating system was Ubuntu 18.04.5 LTS. Disk was fast networked file-systems; a mixture of Lustre and NetApp NFS. We were unable to clear the disk cache for these devices, but the tasks are mainly CPU bound so it is unlikely that disk I/O would become a significant bottleneck.

For CRAM, reference sequences were held as raw sequences (named after their md5sum) and fetched via NFS. Deez references were supplied as FASTA and Genozip used its own compressed reference format.

#### 1 Tool version numbers

**Deez:** v1.9-beta2-26-g92cd56b (github clone)

**SAMtools:** 1.13 (linked against HTSlib 1.13 with libdeflate support).

**HTScodecs:** 1.1.1 Used internally by SAMtools/HTSlib and CRAM benchmarks.

**HTScodecs:** jkbonfield/unroll32 branch (6c5cb8d) used for SIMD tests.

**MPEG-G:** GenomSys encoding library 4.0 (personal communication).

**Genie/GABAC:** Develop branch (0382d5d)

**Genozip:** 12.0.33-4-g308a84dd and 12.0.34-6-g548624e0 (PacBio CLR)

**SNAP:** 1.0beta.24

#### 1.1 Tool command lines

**SAMtools:** Encoding and decoding with SAMtools for BAM and CRAM tests are performed with

`samtools view -@12 -o -T` *ref.fa* `-O` *fmt,options* `-o` *out.fmt in.bam*

`samtools view -@12 -f 0xffff` *in.fmt*

**Deez:** Encoding and decoding with Deez is done via:

`deez -t 12 -!  -v1 -r` *ref.fa in.bam* `-o` *out.dz*

**Genozip:** Encoding and decoding with Genozip uses:

`genozip -e` *ref.genozip* `-@12 -f` *in.bam*

`genocat -e` *ref.genozip* `-@12 -z0 -o /dev/null` *in.bam.genozip*

We did not try the `--optimize` option as the manual states this is a lossy quality value compression.

Scripts to help with benchmarking can be found at https://github.com/jkbonfield/cram31_bench.

### 2 New CRAM 3.1 entropy encoders

CRAM 3.0 has "external" library support via Deflate (bgzf / gzip), Bzip2 and LZMA. It also has it's own static frequency entropy encoder in rANS, using either Order-0 or Order-1 statistics.

CRAM 3.1 modifies the rANS entropy encoder normalisation to be 16-bits at a time for speed (CRAM 3.0 rANS is 8-bit) and adds internal support for bit-packing, run-length encoding and data-striping.

It also adds an alternative adaptive arithmetic encoder, as well as data-specific custom codecs for read name and quality value compression.

Demonstration JavaScript implementations of these codecs also exist in the HTScodecs package, although these should be consider as examples rather than production ready. They do however demonstrate the practicality of implementing CRAM 3.1 in a web environment, with JavaScript speeds typically 5x slower than their C counterpart.

The quality files used in testing here are available for download from ftp://ftp.sanger.ac.uk/pub/users/jkb/CRAM3.1/. Each of these are the first 1 million quality strings from a local NovaSeq (q4+dir) and a HiSeq 2000 run (q40+dir, from 9827_2#49.bam). The first column in each file is the quality string with the second column being a flag for first read (READ1) or second read (READ2). All entropy benchmarks were performed on the first column only with newlines removed, except for FQZComp which utilises the read start points and "selector" value.

#### 2.1 Static 16-bit rANS coding

A modern CPU can dispatch multiple instructions per cycle and permits a degree of out-of-order execution. However there is a delay between starting an operation and the result being available. Hence a series of connected expressions (e.g. $b = a * 2; c = b + 17; d = c - a$) leads to stalling the instruction pipeline.

The existing CRAM 3.0 rANS entropy encoder interleaves 4 rANS states so independent calculations can be performed to avoid pipeline stalling. Each state renormalises 8-bits of data at a time (emitting 8-bits at a time during encoding). This is listed as "rANS4x8" below.

While CRAM 3.1 is still permitted to use the old rANS implementation, it adds a newer version which renormalises with 16-bits at a time for speed. Additionally it can select interleaving either 4 or 32 states. 4 is sufficient to avoid instruction pipelining for traditional scalar maths operations, but with 32 we can use 4 lanes of 8x32-bit SIMD vectors (AVX2) or 2 lanes of 16x32 (AVX512). Unfortunately the compilers are not able to auto-vectorise the code, but we have written multiple SIMD implementations in the HTScodecs github repository with the ability to detect the CPU and execute the fastest available. The CRAM 3.1 rANS codecs are listed as "rANS4x16" and "rANS32x16" below.

We present figures showing the encoding and decoding rates for various rANS options, with and without bit-packing and RLE, and with and varying degrees of vectorisation from none (compiler automatic, mostly traditional scalar code) to AVX512. As we needed AVX512 support, all Intel figures were tested on an Intel Xeon Gold 6242 CPU at 2.8GHz. We also present ARM Neon benchmarks on AWS Neoverse-N1 stepping r3p1.

Tables 1 and 2 show the performance of encoding NovaSeq quality values on Intel and ARM platforms. Huffman encoding is poor due to the need to emit at least 1 bit per symbol. Uncompressed data size is 151MB.

Table 3 shows the encoding of HiSeq 2000 quality values. As with the NovaSeq there is a considerable speed gain available by interleaving 32 rANS states and utilising SIMD, particularly when decoding. The uncompressed data file is 100MB.

In addition to the PACK and RLE methods, rANS also supports a STRIPE filter. This splits the data stream into N sub-streams consisting of the the every $N^{th}$ byte, every $N^{th} + 1$ byte, and so on. Each of these are then compressed using their own rANS parameters. This can be particularly useful for BAM "B" auxiliary arrays. For example an "XX:B:i," auxiliary tag will store 32-bit numbers. If such numbers fall in the 1 to 100,000 range then bits 24-31 will always be 0 and bits 16-23 will be 0 or 1 only, with the bulk of the entropy in the first two bytes per value. STRIPE can permit RLE on the 4th byte, PACK on the 3rd byte, and Order-0 or Order-1 on the first two bytes.

Table 1: NovaSeq quality: entropy encoder speeds. Intel

| Program | Option | Size(MB) | Enc(MB/s) | Dec(MB/s) |
|---|---|---|---|---|
| Zlib | Huffman | 21.48 | 151.3 | 365.4 |
| FSE | tANS | 9.90 | 435.2 | 531.1 |
| FSE | Huffman | 20.74 | 709.8 | 1302.9 |
| rANS4x8 | O0 | 9.91 | 391.4 | 531.1 |
| rANS4x16 | O0 | 9.90 | 446.5 | 823.7 |
| rANS32x16 | O0 | 9.92 | 476.9 | 675.3 |
| rANS32x16-SSE4 | O0 | 9.92 | 476.6 | 1007.5 |
| rANS32x16-AVX2 | O0 | 9.92 | 617.5 | 1609.3 |
| rANS32x16-AVX512 | O0 | 9.92 | 857.0 | 2297.3 |
| rANS4x8 | O1 | 9.14 | 271.2 | 406.8 |
| rANS4x16 | O1 | 9.14 | 285.4 | 551.7 |
| rANS32x16 | O1 | 9.16 | 265.4 | 354.0 |
| rANS32x16-SSE4 | O1 | 9.16 | 264.2 | 675.4 |
| rANS32x16-AVX2 | O1 | 9.16 | 356.7 | 1091.7 |
| rANS32x16-AVX512 | O1 | 9.16 | 476.2 | 1385.5 |
| rANS4x16 | O0,RLE | 10.51 | 426.1 | 905.1 |
| rANS32x16-AVX512 | O0,RLE | 10.54 | 450.6 | 1032.3 |
| rANS4x16 | O0,PACK | 9.10 | 795.6 | 2217.9 |
| rANS32x16-AVX512 | O0,PACK | 9.12 | 896.2 | 3546.8 |
| rANS4x16 | O0,PACK+RLE | 9.15 | 562.6 | 1247.1 |
| rANS32x16-AVX512 | O0,PACK+RLE | 9.18 | 583.6 | 1378.3 |
| rANS4x16 | O1,RLE | 8.74 | 391.8 | 835.4 |
| rANS32x16-AVX512 | O1,RLE | 8.77 | 420.0 | 996.6 |
| rANS4x16 | O1,PACK | 8.47 | 546.3 | 1372.0 |
| rANS32x16-AVX512 | O1,PACK | 8.48 | 607.5 | 1959.0 |
| rANS4x16 | O1,PACK+RLE | 8.24 | 464.4 | 1001.9 |
| rANS32x16-AVX512 | O1,PACK+RLE | 8.26 | 489.3 | 1123.1 |

Block size 1MB, except FSE Huffman which used 128KB.

Table 2: NovaSeq quality: entropy encoder speeds, on Arm NEON

| Program | Option | Size(MB) | Enc(MB/s) | Dec(MB/s) |
|---|---|---|---|---|
| rANS4x8 | O0 | 9.91 | 391.4 | 531.1 |
| rANS4x16 | O0 | 9.90 | 446.5 | 823.7 |
| rANS32x16-Scalar | O0 | 9.92 | 476.9 | 675.3 |
| rANS32x16-Neon | O0 | 9.92 | 476.6 | 1007.5 |
| rANS4x8 | O1 | 9.14 | 271.2 | 406.8 |
| rANS4x16 | O1 | 9.14 | 285.4 | 551.7 |
| rANS32x16-Scalar | O1 | 9.16 | 265.4 | 354.0 |
| rANS32x16-Neon | O1 | 9.16 | 264.2 | 675.4 |

#### 2.2 Adaptive arithmetic coding

The adaptive arithmetic codec is best for non-stationary probabilities where the characteristics of the data may change throughout the block. However this is not a common feature of sequencing data, so the extra complexity is rarely required. This implementation of arithmetic coding is byte-

Table 3: HiSeq2000 quality: Entropy encoder speeds, on Intel

| Program | Option | Size(MB) | Enc(MB/s) | Dec(MB/s) |
|---|---|---|---|---|
| Zlib | Huffman | 50.65 | 116.3 | 257.2 |
| FSE | tANS | 50.12 | 439.9 | 596.1 |
| FSE | Huffman | 50.55 | 721.3 | 1206.3 |
| rANS4x8 | O0 | 50.13 | 350.4 | 523.4 |
| rANS4x16 | O0 | 50.12 | 436.8 | 820.3 |
| rANS32x16 | O0 | 50.13 | 387.1 | 676.4 |
| rANS32x16-SSE4 | O0 | 50.13 | 387.6 | 1004.4 |
| rANS32x16-AVX2 | O0 | 50.13 | 583.6 | 1593.5 |
| rANS32x16-AVX512 | O0 | 50.13 | 795.8 | 2239.0 |
| rANS4x8 | O1 | 48.45 | 241.0 | 353.2 |
| rANS4x16 | O1 | 48.44 | 263.1 | 524.5 |
| rANS32x16 | O1 | 48.45 | 238.7 | 421.4 |
| rANS32x16-SSE4 | O1 | 48.45 | 236.4 | 679.5 |
| rANS32x16-AVX2 | O1 | 48.45 | 325.0 | 998.9 |
| rANS32x16-AVX512 | O1 | 48.45 | 410.4 | 1179.3 |

Block size 1MB, except FSE Huffman which used 128KB.

wise rather than the more usual bit-wise. This can be faster for low-order models, particularly on data sets with only a few symbols being used. As with the CRAM 3.1 rANS implementation, the arithmetic coder also includes the bit-packing, run length encoding and data-striping modes.

For comparisons with the other tools, Tables 4 and 5 show the the same NovaSeq and HiSeq2000 quality data along side a representative non-vectorised rANS line for comparison. Also included is MPEG-G's GABAC algorithm, as implemented in the Genie OpenSource tool. This is also an adaptive arithmetic codec (albeit bit-wise) with the ability to do bit-based transformations before entropy encoding, so is a good comparison against the arithmetic coder used in CRAM 3.1.

Note the Genie gabac-app tool requires little-endian 32-bit data, so the quality data was first transformed with a perl script. This transformation has not been included in the timing figures.

The adaptive arithmetic codec is an order of magnitude slower than rANS, but offers some modest improvement in ratios, particularly the NovaSeq data. GABAC is an order of magnitude slower again, but this is perhaps not indicative of the speed of a fully optimised implementation. It seems poor on compression ratio, but there may also be room for further improvement within the GABAC data format.

Table 4: HiSeq2000 quality: Entropy encoder speeds

| Program | Option | Size(MB) | Enc(MB/s) | Dec(MB/s) |
|---|---|---|---|---|
| rANS4x16 | O0 | 50.12 | 436.8 | 820.3 |
| arith | O0 | 50.05 | 53.4 | 33.4 |
| rANS4x16 | O1 | 48.44 | 263.1 | 524.5 |
| arith | O1 | 48.37 | 45.7 | 31.0 |
| GABAC-app | -d15 -s1 | 50.10 | 3.6 | 5.5 |

Table 5: NovaSeq quality: entropy encoder speeds

| Program | Option | Size(MB) | Enc(MB/s) | Dec(MB/s) |
|---|---|---|---|---|
| rANS4x16 | O0,PACK+RLE | 9.15 | 562.6 | 1247.1 |
| arith | O0 | 9.83 | 120.0 | 94.5 |
| arith | O0,PACK+RLE | 9.16 | 220.0 | 188.3 |
| rANS4x16 | O1,PACK+RLE | 8.24 | 464.4 | 1001.9 |
| arith | O1 | 9.12 | 105.3 | 91.3 |
| arith | O1,PACK+RLE | 8.12 | 156.5 | 115.1 |
| GABAC-app | -d15 -s1 | 8.40 | 5.0 | 8.0 |

#### 2.3 Generalised FQZComp

The original FQZComp tool had a choice of three quality models, selected by parameters -q1, -q2 and -q3 using 12-bits, 16-bits and 20-bits respectively for the model context. The construction of these contexts were hard coded.

CRAM 3.1's FQZComp model is designed around block-based compression offering random access, so uses at most 16-bits of context. The construction of this context is configurable with the construction rules stored in the data stream so the decoder can follow the instructions during decode. This offers considerable freedom in context construction.

For example, the base qualities on long read technologies may not vary much along the length of the read, so attempting to utilise position on the sequence will likely harm compression by making the model slower to adapt. Conversely Illumina data has a strong bias between the 5' and 3' end. Additionally this also means knowing the orientation can help improve compression.

CRAM's codecs are designed to be operated in isolation without knowledge of other CRAM fields. This is a deliberate design decision in order to remove data dependencies and permit a more column-based filtering access pattern. However the additional bit required per read to store (duplicate) the orientation within the quality stream may be more than offset by space saving. It also permits a flag per read to indicate entire quality string duplication, which is beneficial when supplementary or secondary alignments are present.

Other data available for use in context construction are previous quality values, a running total of successive quality differences since the start of this read, and the position along the current read. These values can be indirected via a lookup table, so they do not need to be uniformly distributed.

We can also store part of the context from the data stream itself, stored at the start of each new record. We use this to classify data by a variety of methods. For example separating by READ1 and READ2 flags can sometimes be beneficial. We may also wish to group reads by their average quality value, the number of differences, or even their position on the flow cell (extracted from the read name) with edge-effects or bubbles changing quality characteristics. Any categorisation can be considered by the encoder as the decoder simply uses data from the file format without knowing how it was derived. These are known as the "selector" bits.

Pseudo-code for context construction used during the decoding process is below. See CRAM-codecs specification for more details.

Table 6 shows the performance of FQZComp on the HiSeq 2000 and NovaSeq test sets, with several different context configurations. Also shown is libbsc, as a high quality commonly used general purpose compression tool having a similar data throughput. We also repeat the best rANS and arith ratios for ease of comparison.

```
function FQZUpdateContext(params, q)
    ctx ← params.context                                   ▷ Also the initial value
    qctx ← (qctx << params.qshift) + qtab_q
    ctx ← ctx + ((qctx AND (2^params.qbits − 1)) << params.qloc)
    if params.pflags AND 32 then                           ▷ have_ptab
        p ← Min(pos, 1023)
        ctx ← ctx + (ptab_p << params.ploc)
    end if
    if params.pflags AND 64 then                           ▷ have_dtab
        d ← Min(delta, 255)
        ctx ← ctx + (dtab_d << params.dloc)
        if prevq ≠ q then
            delta ← delta + 1
        end if
        prevq ← q
    end if
    if params.pflags AND 8 then                            ▷ do_sel
        ctx ← ctx + (sel << params.sloc)
    end if
    return ctx AND (2^16 − 1)
end function
```

Table 6: Quality value FQZComp performance

| Program | Option | Size(MB) | Enc(MB/s) | Dec(MB/s) |
|---|---|---|---|---|
| *NovaSeq qualities* | | | | |
| rANS4x16 | O1,PACK+RLE | 8.24 | 464.4 | 1001.9 |
| arith | O1,PACK+RLE | 8.12 | 156.5 | 115.1 |
| bsc | -m0e2tTp | 7.72 | 19.4 | 35.5 |
| FQZComp | -s0 | 7.27 | 28.6 | 51.8 |
| FQZComp | -s1 | 7.29 | 26.9 | 45.7 |
| FQZComp | -s1+strand | 7.21 | 21.4 | 26.7 |
| *HiSeq 2000 qualities* | | | | |
| rANS4x16 | O1 | 48.4 | 263.1 | 524.5 |
| arith | O1 | 48.4 | 45.7 | 31.0 |
| bsc | -m0e2tTp | 43.8 | 6.8 | 9.1 |
| FQZComp | -s0 | 42.5 | 15.8 | 17.8 |
| FQZComp | -s1 | 34.0 | 16.4 | 18.8 |
| FQZComp | -s1+strand | 31.3 | 13.3 | 14.9 |

Compression of NovaSeq and HiSeq 2000 quality values using libbsc and CRAM 3.1's FQZComp, in 10 blocks of 100,000 records.

#### 2.4 Name tokeniser

The name tokeniser deconstructs a read name into a series of elements and then compares each element against a previous tokenised read name. Typically this is the immediately previous name, but with mixed data sets it may be an earlier one matching the same pattern. These tokens are then collated in columns with each column then compressed using either the CRAM 3.1 rANS or adaptive arithmetic coders.

Table 7 shows the compression performance on position-sorted and name-sorted data files. We show results for our name tokeniser (tok3) against brotli (bro), libbsc (bsc), bzip2, gzip (libdeflate implementation), mcm, xz and zstd.

The position sorted data is the aligned NovaSeq file used in the CRAM benchmarks (ERR3239334). The read name sorted data set is the interleaved read1/read2 records from ERR174326. Both data sets are 1 million names, separated into 10 lots of 100,000 names to simulate coarse random access capabilities.

Mcm and libbsc are strong contenders, with mcm sometimes ahead of the CRAM name tokeniser on ratio, however as with other codecs CRAM aims to strike the balance between speed and size.

Table 7: Name / identifier compression

| | Name sorted | | | Position sorted | | |
|---|---|---|---|---|---|---|
| Program | Size | Enc | Dec | Size | Enc | Dec |
| tok3 -3 | 1.11 | 24.68 | 49.61 | 3.55 | 22.98 | 67.84 |
| tok3 -7 | 0.89 | 15.46 | 48.54 | 3.55 | 10.00 | 69.44 |
| tok3 -19 | 0.88 | 8.82 | 41.59 | 3.48 | 4.84 | 39.13 |
| bro -3 | 2.90 | 90.49 | 164.71 | 7.03 | 67.13 | 176.46 |
| bro -9 | 2.79 | 9.68 | 177.69 | 5.91 | 6.28 | 203.53 |
| bro -11 | 1.51 | 0.47 | 164.71 | 5.01 | 0.36 | 183.18 |
| bsc -m0e2tT | 2.60 | 8.27 | 29.29 | 4.06 | 7.50 | 28.00 |
| bsc -m5e1tT | 1.68 | 27.61 | 23.07 | 3.99 | 22.59 | 19.53 |
| bzip2 | 2.52 | 32.08 | 222.11 | 6.18 | 15.97 | 137.88 |
| gzip -6 | 2.87 | 75.76 | 254.95 | 6.58 | 54.95 | 240.42 |
| gzip -12 | 2.14 | 3.78 | 250.59 | 5.50 | 2.80 | 241.93 |
| mcm -t1 | 1.42 | 5.31 | 5.57 | 3.59 | 4.73 | 4.80 |
| mcm -m7 | 1.22 | 3.07 | 3.21 | 3.43 | 2.69 | 2.78 |
| xz -1 | 2.15 | 27.32 | 78.81 | 6.53 | 15.84 | 61.06 |
| xz -9 | 1.31 | 2.50 | 74.98 | 4.87 | 1.70 | 65.31 |
| zstd -9 | 2.51 | 39.73 | 229.05 | 6.22 | 21.49 | 236.00 |
| zstd -19 | 1.56 | 2.61 | 225.53 | 5.02 | 1.81 | 224.96 |

Size in MB. Encode and Decode rates in MB/s

### 3 Detailed CRAM file compression results: aligned data

The "io_lib" `cram_dump` tool can be used to interrogate the layout of a CRAM file. An example container header is listed below.

```
Container_header block pos 4628
  Preservation map:
    RN => 1 (0x1)
    SM => 94095715340138 (0x55945db00f6a)
    TD => 94095715340100 (0x55945db00f44)
    AP => 0 (0x0)
  Substitution map:
    A: CGTN
    C: AGTN
    G: ACTN
    T: ACGN
    N: ACGT
  TD map:
      0: PLZPUZLBZSMZPGZ
      1: PLZPUZLBZSMZPGZNMC

  Record encoding map:
     FN =>         EXTERNAL {26}
     FP =>         EXTERNAL {28}
     RL =>          HUFFMAN {1, 101, 1, 0}
     RN =>  BYTE_ARRAY_STOP {0, 11}
     QS =>         EXTERNAL {12}
     BA =>         EXTERNAL {30}
     BB =>   BYTE_ARRAY_LEN {1, 1, 42, 1, 1, 37}
     TL =>         EXTERNAL {32}
     IN =>  BYTE_ARRAY_STOP {0, 13}
     BF =>         EXTERNAL {15}
     MF =>          HUFFMAN {1, 0, 1, 0}
     TS =>          HUFFMAN {1, 0, 1, 0}
     CF =>          HUFFMAN {1, 3, 1, 0}
     AP =>             BETA {0, 28}
     FC =>         EXTERNAL {27}
     MQ =>          HUFFMAN {1, 0, 1, 0}
     DL =>         EXTERNAL {29}
     BS =>         EXTERNAL {31}
     NP =>          HUFFMAN {1, 0, 1, 0}
     SC =>  BYTE_ARRAY_STOP {0, 14}
     NS =>          HUFFMAN {1, 255, 255, 255, 255, 15, 1, 0}
     RG =>          HUFFMAN {1, 0, 1, 0}
     RI =>         EXTERNAL {33}

  Tag encoding map:
    PUZ =>  BYTE_ARRAY_STOP {9, 224, 80, 85, 90}
    LBZ =>  BYTE_ARRAY_STOP {9, 224, 76, 66, 90}
    SMZ =>  BYTE_ARRAY_STOP {9, 224, 83, 77, 90}
    NMC =>   BYTE_ARRAY_LEN {3, 4, 1, 1, 1, 0, 1, 4, 224, 78, 77, 67}
    PGZ =>  BYTE_ARRAY_STOP {9, 224, 80, 71, 90}
    PLZ =>  BYTE_ARRAY_STOP {9, 224, 80, 76, 90}
```

The record encoding map describes how each data series is encoded. This is mostly outputting to blocks as-is (EXTERNAL) or with variable length data (either null terminated or with explicit length encodings). HUFFMAN is also (ab)used in this example for constant items, with a Huffman tree containing a single zero-bit long symbol. The BETA encoding is also used for alignment positions. The parameters to each encoding describe, among other things, the block it is stored in, for example soft-clipped sequence ("SC") is written to block number 14.

The blocks are then grouped together in slices, and each block describes the compression codec used for that data type.

```
Slice 1/1, container offset 294, file offset 4922
    Slice content type MAPPED_SLICE
    Slice ref seq    -2
    Slice ref start  0
    Slice ref span   0
    Slice MD5        00000000000000000000000000000000
    Rec counter      0
    No. records      25000
    No. blocks       20
    Blk IDS:         {11, 12, 13, 14, 15, 26, 27, 28, 29, 30, 31, 32, 33, 5261146, 5131587, 5264730, 5459290, 4997722, 5262426}
    Ref base id:     -1
```

```
Block 1/20
    Size:         86843 comp / 87500 uncomp
    Method:       GZIP  (1)
    Content type: CORE
    Content id:   0

Block 2/20
    Size:         24561 comp / 746680 uncomp
    Method:       TOK3_R        (8)
    Content type: EXTERNAL
    Content id:   11
    Keys:         RN

Block 3/20
    Size:         676771 comp / 2525000 uncomp
    Method:       FQZ   (7)
    Content type: EXTERNAL
    Content id:   12
    Keys:         QS
```

The "io_lib" `cram_size tool` can summarise all this data, reporting how much space each data series consumes and which compression methods are used (this may be more than one as a different codecs may be used in different slices).

```
Block CORE              , total size  1439857150
Block content_id      11, total size   403313878               n            RN
Block content_id      12, total size 12771259069              f             QS
Block content_id      13, total size     2632067      01                    IN
Block content_id      14, total size   194147009       1  8                 SC
Block content_id      15, total size    50899663       1  8                 BF
Block content_id      26, total size    80976004      01                    FN
Block content_id      27, total size    38717234      01  8                 FC
Block content_id      28, total size   219404316 g    0                     FP
Block content_id      29, total size     1517445      0                     DL
Block content_id      30, total size   282950932       1   9                BA
Block content_id      31, total size    47329198      014                   BS
Block content_id      32, total size     9824542      01                    TL
Block content_id      33, total size   185392546  b      5                  RI
Block content_id 4997722, total size      332138            2               LBZ
Block content_id 5131587, total size     3146066      0                     NMC
Block content_id 5261146, total size     4350689 g         9                PGZ
Block content_id 5262426, total size     1228921           9                PLZ
Block content_id 5264730, total size      332138            2               PUZ
Block content_id 5459290, total size      332256 g          2               SMZ
```

In the tables below, we report total file size and time taken to encode and decode, gathered using the `time` command. This shows the difference between total CPU usage across all threads and the elapsed time, giving an indication of thread scalability. The entry for "Deez q1" corresponds to the `-q1` option of Deez which uses the samcomp quality model. Random access is not available on the quality field for this method, so it is excluded from the main figure.

We also include numbers for the experimental CRAM 4.0 version. This is ongoing work and is expected to change further, but the primary reason for size reduction over 3.1 is deduplication of read names and the option to store quality values in their original 5' to 3' orientation.

Figure 1 is a zoomed up colourised version of the primary figure in the paper, showing more details on the relative performance of the different CRAM versions. Because of the zooming, BAM is off the top of the chart and Deez's poor multi-threading scaling mostly hides the results to the right. Figure 2 is the same data showing the CPU time (User + System). This now includes Deez, which is quite performant. MPEG-G is not shown as we only have the elapsed time for that test system.

Encode and Decode timings: Elapsed time (12 threads)

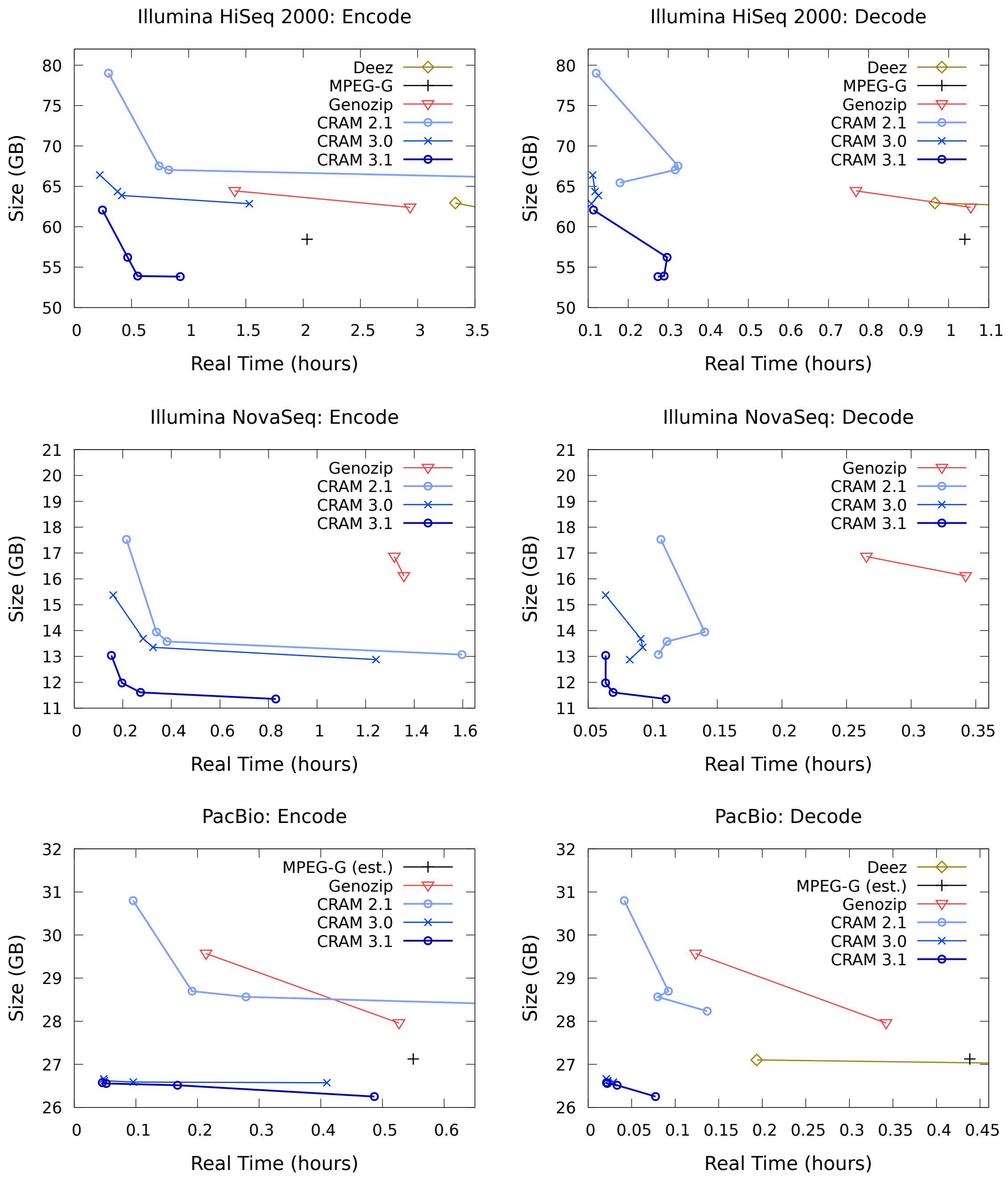


Figure 1: Multi-threaded benchmark of aligned data formats showing the elapsed time. MPEG-G figures are taken from the Voges et.al paper, with "MPEG-G (est.)" assumed to be working on the same data file as our CRAM measurements, but this is not certain. (See text.) Deez is absent as the elapsed time is too large. BAM is not shown as the file sizes are too big.

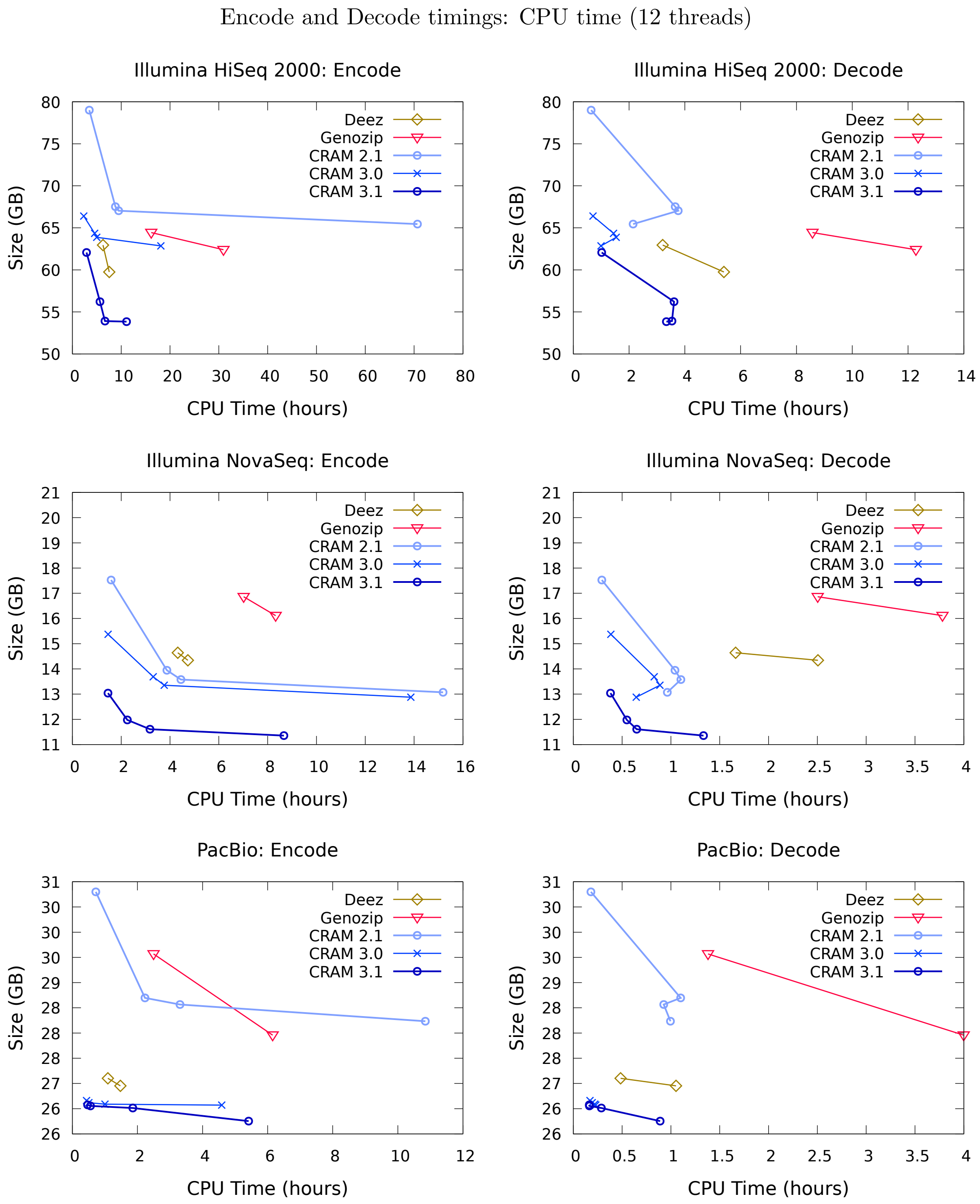


Figure 2: Multi-threaded benchmarks of aligned data formats showing the CPU (User + System) time. MPEG-G figures are taken from the Voges et.al paper, with "MPEG-G (est.)" assumed to be working on the same data file as our CRAM measurements, but this is not certain. (See text.) BAM is not shown as the file sizes are too big. MPEG-G is absent as only elapsed time was reported.

#### 3.1 NovaSeq data

We tried to obtain details on how the MPEG-G dataset 37 was aigned, in order to compare tools against MPEG-G, but were unable to obtain them or the BAM file. Hence the results below use a difference public data set.

Source: ftp://ftp.sra.ebi.ac.uk/vol1/run/ERR323/ERR3239334/NA12878.final.cram

Figure 3 shows the break down in data type sizes for CRAM 2.1 (normal and small profiles), CRAM 3.0 (normal, small and archive level 9), CRAM 3.1 (normal, small and archive level 9), CRAM 4.0 (normal and small), Deez (–q1 and –q2) and Genozip (normal and –fast). Sequence size is everything that is not Name, Qual or Auxiliary fields and hence also includes mapping quality, template lengths and flags. This is to make it easier to compare between tools.

Deez is significantly ahead on sequence here, presumably due to the two-level delta, while the new CRAM codecs are significantly ahead on read names and quality values.

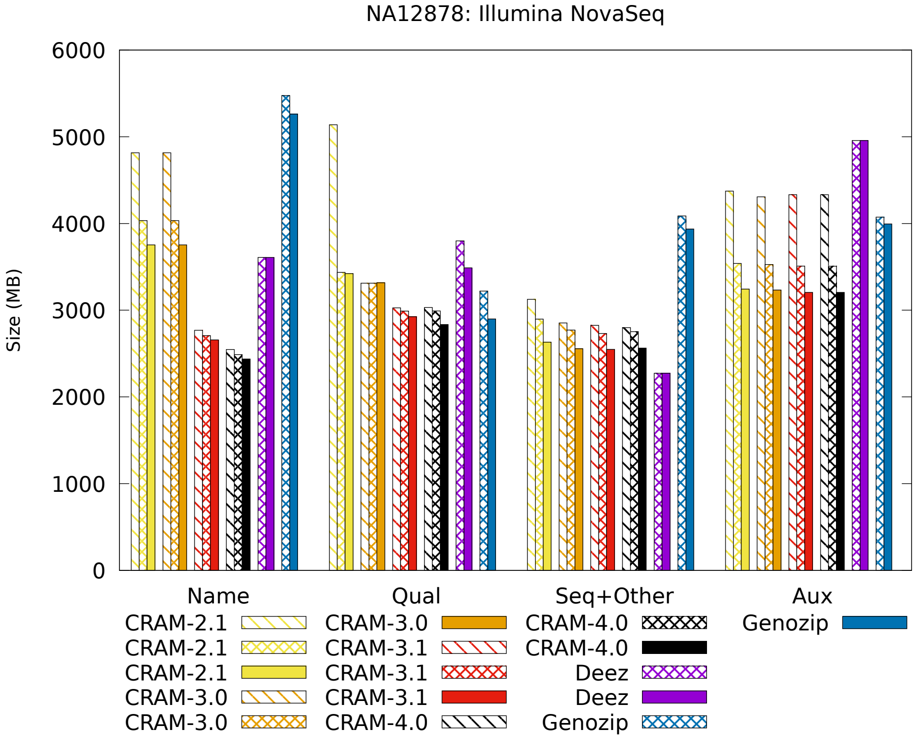


Figure 3: A break down of the relative size for each data type. Successive bars for the same format represent more advanced compression options.

Table 8: NovaSeq detailed compression timings

| Format | Size (bytes) | Encode time(s) CPU | Encode time(s) Elapsed | Decode time(s) CPU | Decode time(s) Elapsed |
|---|---|---|---|---|---|
| BAM normal | 42,157,574,307 | 5,134 | 703 | 767 | 125 |
| BAM small | 40,552,289,097 | 39,145 | 3,948 | 766 | 125 |
| BAM archive | 39,457,586,755 | 123,021 | 10,360 | 767 | 129 |
| CRAM 2.1 normal | 17,527,458,203 | 5,710 | 774 | 1,050 | 383 |
| CRAM 2.1 small | 13,944,513,714 | 13,931 | 1,218 | 3,750 | 505 |
| CRAM 2.1 archive | 13,579,314,010 | 16,002 | 1,376 | 3,966 | 400 |
| CRAM 2.1 archive9 | 13,073,007,080 | 54,670 | 5,750 | 3,467 | 376 |
| CRAM 3.0 normal | 15,373,623,293 | 5,276 | 576 | 1,385 | 229 |
| CRAM 3.0 small | 13,685,309,202 | 11,913 | 1,022 | 2,986 | 327 |
| CRAM 3.0 archive | 13,351,949,900 | 13,540 | 1,167 | 3,195 | 332 |
| CRAM 3.0 archive9 | 12,876,075,348 | 49,919 | 4,475 | 2,321 | 296 |
| CRAM 3.1 normal | 13,038,275,610 | 5,254 | 551 | 1,374 | 229 |
| CRAM 3.1 small | 11,977,349,891 | 8,108 | 708 | 1,973 | 229 |
| CRAM 3.1 archive | 11,609,660,736 | 11,442 | 982 | 2,339 | 250 |
| CRAM 3.1 archive9 | 11,356,337,356 | 31,187 | 2,988 | 4,800 | 397 |
| CRAM 4.0 normal | 12,799,631,141 | 4,962 | 491 | 1,404 | 256 |
| CRAM 4.0 small | 11,779,546,512 | 7,923 | 684 | 2,043 | 318 |
| CRAM 4.0 archive | 11,267,211,530 | 14,378 | 1,213 | 4,822 | 402 |
| CRAM 4.0 archive9 | 11,046,703,345 | 29,174 | 2,801 | 4,829 | 441 |
| Deez (normal) | 14,641,046,003 | 15,552 | 9,119 | 5,984 | 1,479 |
| Deez q2 | 14,341,960,807 | 17,039 | 9,994 | 9,022 | 3,929 |
| Deez q1 | 13,840,305,198 | 17,427 | 10,314 | 9,736 | 4,505 |
| Genozip fast | 16,862,262,618 | 25,267 | 4,746 | 9,012 | 955 |
| Genozip (normal) | 16,114,441,352 | 29,977 | 4,887 | 13,613 | 1,232 |

Note: Deez -q1 lacks random access for quality values.

#### 3.2 HiSeq 2000 data

This is the Illumina Platinum Genomes deep sequencing of NA12878. It is also the MPEG-G Data-Set 02.

Source: ftp://ftp.sra.ebi.ac.uk/vol1/run/ERR194/ERR194147/NA12878_S1.bam

The MPEG-G format quoted size from their paper supplementary material is 58444073745 bytes, at 8,280s to encode and 3,990s to decode using 8 threads on an Intel i7-7700 running at 3.6Hz. The earlier MPEG document M56361 quotes the size as a very similar 58457106765, taking 7,320s to encode and 3,745s to decode using 12 threads on an Intel E5-2670 running at 2.6GHz. Our tests are on an Intel E5-2660, also with 12 threads, nominally running at 2.2GHz but with CPU frequency scaling potentially overclocking this. We believe the speed benchmarks to be sufficiently similar to be of value in comparisons, although we cannot quantify the impact of any I/O speed differences.

Figure 4 shows the break down in data type sizes for CRAM 2.1 (normal and small profiles), CRAM 3.0 (normal, small and archive level 9), CRAM 3.1 (normal, small and archive level 9), Deez (–q1 and –q2), MPEG-G and Genozip (normal and –fast). Sequence size is everything that is not Name, Qual or Auxiliary fields and hence also includes mapping quality, template lengths and flags. This is to make it easier to compare between tools.

The quality values dominate the file size here, giving CRAM 3.1 a significant advantage in the overall file size. However the new name takeniser is clearly a visible gain too.

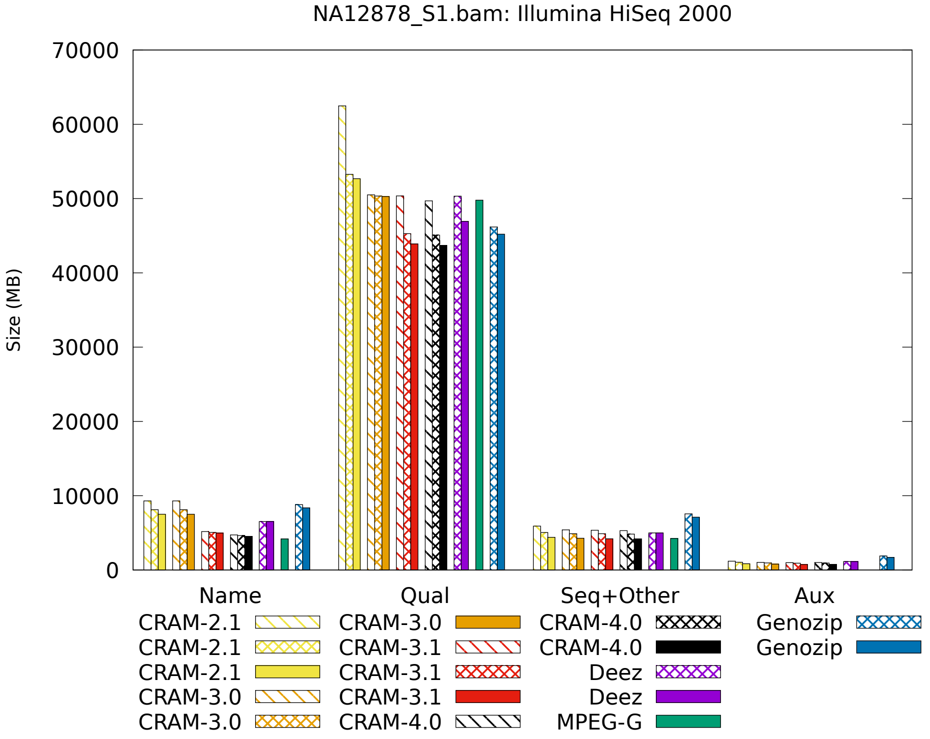


Figure 4: A break down of the relative size for each data type. Successive bars for the same format represent more advanced compression options.

Table 9: HiSeq 2000 detailed compression timings

| Format | Size (bytes) | Encode time(s) CPU | Encode time(s) Elapsed | Decode time(s) CPU | Decode time(s) Elapsed |
|---|---|---|---|---|---|
| BAM normal | 122,405,272,273 | 11,373 | 1,529 | 1,590 | 236 |
| BAM small | 117,643,793,672 | 71,905 | 6,479 | 1,594 | 289 |
| BAM archive | 116,251,338,963 | 135,377 | 11,464 | 1,584 | 265 |
| CRAM 2.1 normal | 79,011,532,839 | 12,502 | 1,078 | 2,310 | 433 |
| CRAM 2.1 small | 67,523,048,483 | 31,712 | 2,668 | 13,155 | 1,168 |
| CRAM 2.1 archive | 67,031,004,053 | 34,091 | 2,970 | 13,541 | 1,142 |
| CRAM 2.1 archive9 | 65,446,690,950 | 254,456 | 21,427 | 7,725 | 646 |
| CRAM 3.0 normal | 66,403,160,848 | 8,338 | 803 | 2,539 | 401 |
| CRAM 3.0 small | 64,351,062,035 | 16,394 | 1,361 | 5,197 | 423 |
| CRAM 3.0 archive | 63,877,044,117 | 17,975 | 1,492 | 5,540 | 453 |
| CRAM 3.0 archive9 | 62,859,694,562 | 65,206 | 5,506 | 3,575 | 390 |
| CRAM 3.1 normal | 62,069,701,318 | 10,391 | 887 | 3,662 | 408 |
| CRAM 3.1 small | 56,209,721,947 | 20,318 | 1,679 | 13,009 | 1,070 |
| CRAM 3.1 archive | 53,909,057,310 | 24,064 | 1,992 | 12,737 | 1,043 |
| CRAM 3.1 archive9 | 53,832,706,189 | 39,905 | 3,334 | 12,013 | 987 |
| CRAM 4.0 normal | 60,899,248,397 | 9,746 | 836 | 3,580 | 455 |
| CRAM 4.0 small | 55,539,803,675 | 19,537 | 2,023 | 12,781 | 1,051 |
| CRAM 4.0 archive | 53,677,139,774 | 22,238 | 1,843 | 11,651 | 949 |
| CRAM 4.0 archive9 | 53,144,998,170 | 42,953 | 3,627 | 11,776 | 964 |
| Deez (normal) | 62,950,633,129 | 22,563 | 11,988 | 11,545 | 3,477 |
| Deez q2 | 59,751,295,736 | 27,163 | 16,077 | 19,436 | 10,434 |
| Deez q1 | 54,883,099,246 | 29,820 | 18,743 | 22,564 | 13,495 |
| Genozip fast | 64,441,271,578 | 58,137 | 5,047 | 30,870 | 2,767 |
| Genozip (normal) | 62,396,471,693 | 111,425 | 10,560 | 44,227 | 3,799 |
| MPEG-G (estimated) | 58,444,073,745 | N/A | *8,280 | N/A | *3,990 |

Note: Deez -q1 lacks random access for quality values.
(*)MPEG-G timings are for a different system. See text for details.

##### 3.3 PacBio CLR data

Source: ftp://ftp.1000genomes.ebi.ac.uk/vol1/ftp/technical/working/20131209_na12878_pacbio/si/NA12878.pacbio.bwa-sw.20140202.bam

The document "Results_MPEG-G_Detailed_M56361.xlsx" states 13,766,776 records, The original BAM file has 25,968,256 records with 13,682,101 primary alignments and 12,286,155 secondary alignments. Given that secondary alignments share the same sequence and quality as their primary, it may be possible to deduplicate these. MPEG-G merges records from the same template together if they occur within the same Access Unit, which is stated to be blocks of 65,536 records. 68% of the secondary alignments are within 65,536 of their primary, so this cannot account for the disparity in the number of records. This document also states "BAM AUX: NM, MD, RG".

Dropping all secondary alignments and most auxiliary tags with `samtools view -F 0x100 -x AS -x PG -x SA -x XS` produces a BAM file closely matching the one quoted in M56361 (it is 0.7% smaller, but that may be due to Deflate implementation differences) and an even closer match to the CRAM 3.0 size (0.3% smaller). We attempted to clarify this, but received no answer. Therefore we are working on the assumption this transformation was applied and have done the same on our results for purposes of a fair comparison. Our results are valid between the tests we performed on versions of CRAM, Deez and Genozip, but we have listed MPEG-G sizes as "estimated" due to this lack of certainty.

Unlike the HiSeq data, timings for the MPEG-G results come from the MPEG-G paper supplementary results and not the M56361 publication. Note these were performed using 8 threads on an Intel Core i7-7700 CPU running at 3.6 GHz. This is considerably different to our results using 12 cores on an Intel Xeon E5-2660 (nominally running at 2.2 GHz but with CPU frequency scaling sometimes temporarily overclocking this). However we note the HiSeq data set has speeds reported on both the i7-7700 system and the E5-2660 with encode times of 8,280 and 7,320 respectively and decode times of 3,990 and 3,745 respectively. This implies the MPEG-G timings below may be between 7 and 12% speed higher than if they were executed on the E5-2660 system. Although the MPEG-G timings are not directly comparable, we believe them to be broadly in the same ballpark and certainly within $\pm 50\%$.

Figure 5 shows the break down in data type sizes. As the sequences are long, read names are a tiny proportion of the total data. The PacBio qualities are dominant and hard to compress effectively. As explained above auxiliary tags were removed, but Genozip (default mode) still stored the MD and NM tags. As in the HiSeq test, the data type sizes for MPEG-G come from the M56361 document. Although this was an earlier report, the total file size matches the one listed in their supplementary results so we believe them to be valid.

We also repeated the PacBio CRAM 3.1 test on a different system with AVX512 support to test the fastest rANS implementations. This test was with 4 cores on an Intel Xeon Gold 6142 CPU (nominally at 2.6GHZ but with auto-scaling enabled). The reduced core count is to ensure a long enough test and to avoid I/O effects. Results averaging over 3 runs are presented in Table 11.

Time saved varies by data set and codec being used. This file has about 77% of file contents in the quality data series, so a SIMD order-1 rANS codec makes a significant difference (16 to 22%). However the slower archival compression profiles would not save nearly as much, nor will files containing a lot of long textual auxiliary tags (such as "XA:Z:"). A small sample of NovaSeq data showed a modest 4% speed gain. It was also observed that using many threads reduced the impact as other factors become more dominant. Furthermore, without CPU pinning the clock speed of CPUs running AVX512 was generally lower, reducing the potential gains.

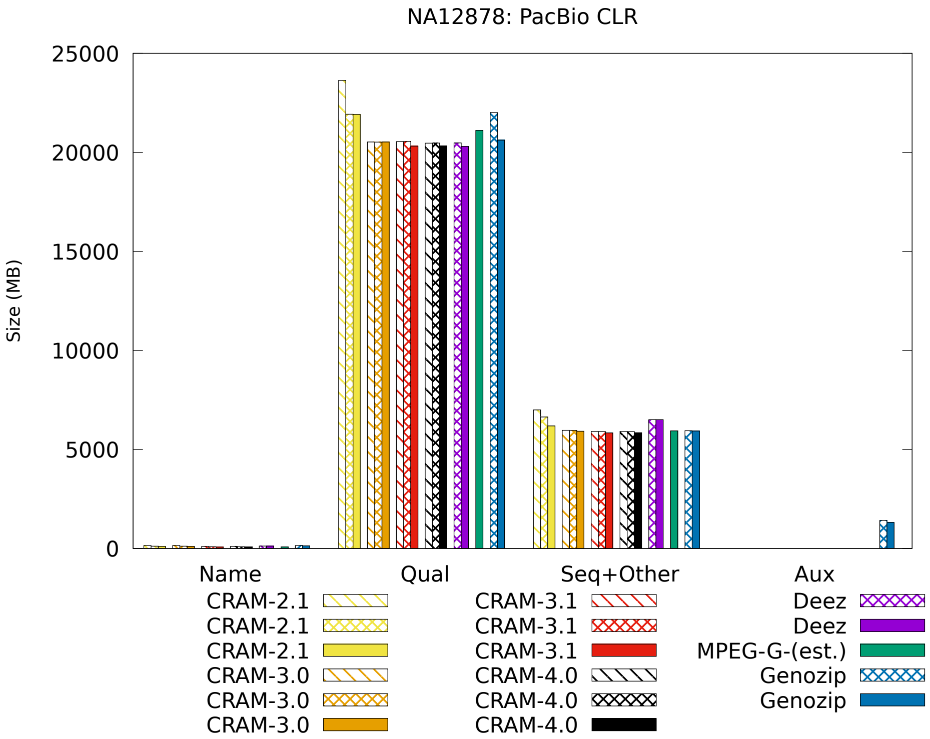


Figure 5: A break down of the relative size for each data type. Successive bars for the same format represent more advanced compression options.

Table 10: PacBio CLR detailed compression timings

| Format | Size (bytes) | Encode time(s) | | Decode time(s) | |
|---|---|---|---|---|---|
| | | CPU | Elapsed | CPU | Elapsed |
| BAM normal | 45,157,801,387 | 2,661 | 483 | 524 | 72 |
| BAM small | 43,195,408,084 | 10,441 | 1,056 | 516 | 69 |
| BAM archive | 42,955,761,594 | 14,706 | 1,357 | 520 | 63 |
| CRAM 2.1 normal | 30,798,998,010 | 2,585 | 344 | 652 | 149 |
| CRAM 2.1 small | 28,696,570,578 | 8,031 | 688 | 3,959 | 332 |
| CRAM 2.1 archive | 28,564,789,419 | 11,908 | 1,003 | 3,337 | 288 |
| CRAM 2.1 archive9 | 28,234,710,246 | 39,049 | 3,982 | 3,583 | 492 |
| CRAM 3.0 normal | 26,661,310,803 | 1,559 | 172 | 615 | 75 |
| CRAM 3.0 small | 26,618,566,639 | 1,822 | 176 | 697 | 81 |
| CRAM 3.0 archive | 26,588,901,633 | 3,604 | 344 | 792 | 104 |
| CRAM 3.0 archive9 | 26,570,268,123 | 16,525 | 1,475 | 830 | 103 |
| CRAM 3.1 normal | 26,575,440,966 | 1,686 | 164 | 588 | 76 |
| CRAM 3.1 small | 26,555,118,552 | 1,995 | 187 | 592 | 79 |
| CRAM 3.1 archive | 26,512,672,102 | 6,681 | 603 | 1,036 | 120 |
| CRAM 3.1 archive9 | 26,252,360,986 | 19,502 | 1,753 | 3,201 | 279 |
| CRAM 4.0 normal | 26,479,249,152 | 1,709 | 189 | 606 | 77 |
| CRAM 4.0 small | 26,456,950,989 | 2,015 | 186 | 610 | 79 |
| CRAM 4.0 archive | 26,424,189,720 | 6,605 | 621 | 1,036 | 117 |
| CRAM 4.0 archive9 | 26,235,991,537 | 20,432 | 1,797 | 3,155 | 272 |
| Deez (normal) | 27,101,199,739 | 3,925 | 2,917 | 1,746 | 697 |
| Deez q2 | 26,952,068,252 | 5,313 | 3,571 | 3,788 | 2,670 |
| Deez q1 | 26,918,516,745 | 5,680 | 3,934 | 4,241 | 3,128 |
| Genozip fast | 29,569,274,634 | 8,983 | 770 | 4,967 | 444 |
| Genozip (normal) | 27,957,030,849 | 22,148 | 1,897 | 14,397 | 1,232 |
| MPEG-G (estimated) | 27,124,417,537 | N/A | *1,980 | N/A | *1,578 |

Note: Deez -q1 lacks random access for quality values.
(*)MPEG-G timings are for a different system. See text for details.

Table 11: PacBio CLR AVX512 compression timings, using 4 threads

| Format | Size (bytes) | Encode time(s) | | Decode time(s) | |
|---|---|---|---|---|---|
| | | CPU | Elapsed | CPU | Elapsed |
| CRAM 3.1 Scalar | 26,574,431,640 | 1,044.8 | 274.8 | 380.7 | 96.5 |
| CRAM 3.1 AVX512 | 26,578,918,156 | 858.4 | 236.1 | 319.7 | 82.0 |

### 4 CRAM compression: unaligned data

Sequencing data is typically generated in FASTQ. We would not consider this to be a suitable long term storage format as aligning or assembling the data both improves compression ratios and also substantially boosts the usefulness. However as an interim solution, data compression is still valuable.

Traditionally standard text compressors (such as gzip) have been used. There are a wide variety of dedicated FASTQ compressors, but we limit ourselves to evaluating the same multi-purpose encoders listed above with unaligned data.

The most naive solution is simply to generate an unaligned BAM, CRAM and related formats. This is a fast workable solution and better than straight FASTQ as it permits the presence of sample meta-data in the file header and has a rigorously defined way to attach additional per sequence meta-data, such as bar-coding, instead of exploiting the FASTQ comment section to create a format within a format.

Our data set is ERR174310_1.fastq.gz and ERR174310_2.fastq.gz from the MPEG-G data set 01-1.

The ERR174310_1.fastq.gz file is in NCBI SRA format, which unfortunately is a non-standard FASTQ variant where the original read identifier is in column 2. To correct this a fix the `samtools import` was made in PR#1485. The command to convert from fastq pairs to BAM is `samtools import -@8 -N -1 ERR174310_1.fastq.gz -2 ERR174310_2.fastq.gz -o ERR174310.u.bam`. The `samtools fastq` command can be used to reverse the process.

Table 12 shows encoding performance for various tools. The MPEG-G line here comes from their paper, so was on an Intel i7-7700. All others were benchmarked by us on an Intel Xeon Gold 6142 using 8 threads. This is a different system to the aligned data benchmarks above as the aligner needed more memory. Note `samtools import` currently has no multi-threaded decoding of FASTQ, so threading scalability is limited. However this is a tool chain implementation issue rather than a limitation of file formats so we expect this to improve.

Table 12: Compression of unaligned FASTQ

| Format | Size (bytes) | CPU(s) | Elapsed(s) |
|---|---|---|---|
| FASTQ.bgzf | 36,770,864,911 | 3,534 | 1,017 |
| BAM | 35,141,235,517 | 2,857 | 1,184 |
| CRAM 3.0 | 25,310,280,282 | 1,379 | 701 |
| CRAM 3.1 | 24,454,192,729 | 1,857 | 704 |
| CRAM 3.1 small | 23,230,961,730 | 3,883 | 725 |
| MPEG-G no-ref | 24,111,913,018 | 6,166 | 1,193 |
| Genozip | 22,930,340,845 | 11,182 | 1,419 |

MPEG-G timings taken from the Supplementary Data for their paper.
Genozip command line: `genozip -@8 ERR174310_[12].fastq.gz`.

It can be seen that storing FASTQ in unaligned BAM offers little benefit over gzip / bgzf, but using CRAM is a significant improvement due to the columnar nature of the format. This is further improved by the new codecs included in CRAM 3.1. Despite the poor multi-threading scalability of `samtools import`, it is competitive on speed. CRAM 3.1 was only slightly beaten by Genozip on size but ran in a fraction of the time. The default block size for Genozip is 16MB which corresponds to about 65,000 records. This is also comparable to the quoted block size for MPEG-G. CRAM 3.1 defaults to 10,000 short-read records for the normal profile, 25,000 for small profile and 100,000 for archive (not tested above).

A more advanced solution is to perform either a rapid sequence assembly, as originally proposed by Fritz *et al.* or use an existing known reference to do reference based alignment. Such methods do not need to be biologically rigorous as we are not attempting to get the best assembly or the best alignments, but simply to reduce the data volumes instead. There are many dedicated

FASTQ compressors using such techniques, but we limit our comparison to the same tools above that also support aligned data, with bgzf (parallel gzip) as a baseline.

Alignment / assembly considerably reduces the data size, but comes with an additional CPU cost. CRAM does not prescribe a specific method and SAMtools / HTSlib do not have this built in, however we can use very fast aligners for this purpose. (It should be noted that spending too much time on this strategy simply implies you should be converting to fully aligned BAM/CRAM anyway in order to leverage the utility of aligning or assembly rather than purely as a compression method.)

For this research we chose the SNAP aligner. SNAP also cannot cope with the non-standard SRA / ENA FASTQ variant and we also get better CPU performance if we naively align the data as interleaved single-ended data rather than pairs. As a pipeline we use `samtools import` to convert from two FASTQ files to uncompressed unaligned BAM, which is fed into SNAP aligner. The aligner outputs uncompressed aligned SAM, which is finally converted to CRAM. As we don't need fine grained random access on FASTQ, we also increased the number of sequences per CRAM slice to 100,000. We provide a crude `run_aligned.sh` script to demonstrate this process.

The SNAP aligner parameters have been chosen to optimise the speed, as we are not attempting to get good alignments, rather to improve compression ratios. Options were `snap-aligner single grch37.snap -t 8 --b -d 8 -D 0 -f -bam -map -o -sam -`. We used SNAP 1.0beta.24 as this gave better speed and size than the latest release.

Note this is very much a proof of principle as the resulting CRAM cannot be trivially converted back to FASTQ without some processing to split apart the two reads into their /1 and /2 components (given we aligned it without the "paired" option). However sufficient data exists to do this without data loss. The data has also been reordered slightly in large blocks due to the multi-threaded implementation in SNAP. This could be fixed without affecting the size and likely having a minimal impact on speed, but this work has not been done.

The ideal solution would be an aligner which can go directly from compressed FASTQ to compressed CRAM in a single process while retaining the exact file ordering.

Table 13: Compression of unaligned FASTQ

| Format | Size (bytes) | CPU(s) | Elapsed(s) |
|---|---:|---:|---:|
| FASTQ.bgzf | 36,770,864,911 | 3,534 | 1,017 |
| SNAP + BAM | 41,296,355,569 | 11,701 | 1,481 |
| SNAP + CRAM 3.0 normal | 18,023,168,960 | 7,885 | 1,139 |
| SNAP + CRAM 3.0 small 100k | 15,290,788,535 | 9,845 | 1,309 |
| SNAP + CRAM 3.1 normal | 17,105,337,821 | 8,507 | 1,196 |
| SNAP + CRAM 3.1 small 100k | 15,290,788,535 | 10,974 | 1,431 |
| MPEG-G global assembly | 16,053,119,238 | N/A | 5,820 |
| Genozip with ref | 15,061,744,296 | 17,868 | 3,842 |

MPEG-G timings taken from the Supplementary Data for their paper

Table 13 shows alignment and assembly based methods of FASTQ compression. The BAM file is larger than the unaligned BAM, due to storing chromosome name and position, mate-pairing information and auxiliary tags, without any benefits from reference based compression. CRAM shows a considerable improvement in compression ratios, with an expected elevated CPU cost. Note Genozip was ran using options `-@8 --pair --reference h38.snap` while SNAP was running on h37. Genozip gave the smallest file, but was considerably slower than CRAM and the same comments in the unaligned data test regarding block sizes apply.

Using SNAP + CRAM in this way is not the only combination and it is not a production-ready solution yet, however the results demonstrate that the concept of marrying a rapid approximate aligner to CRAM is a viable and performant solution without needing to change to a different encoding format for FASTQ data.

### 5 CRAM compression: Crumble lossy output

The SynDip synthetic diploid genome consisting of CHM1 and CHM13 samples has previously been evaluated for lossy-compression with the Crumble tool.

Unlike many tools, Crumble does not utilise its own file format. Rather it modifies the quality string via a simple experiment: if removing the quality values at a specific loci (after alignment) does not change the variant call and does not significantly alter the variant confidence, then these values can be set to a constant high value if they confirm the call or a constant low value if they disagree. The impact of this is a heavily skewed distribution of quality values with much greater predictability and compression ratios.

To evaluate CRAM 3.1 we compared CRAM 3.0 and CRAM 3.1 on the pre-crumble (original) BAM and the post-crumble BAM. These are shown in Tables 14 and 15. The data was Chromosome 1 of CHM1_CHM13_2.bam from ENA accession PRJEB13208.

Table 14: SynDip compression, pre-Crumble

| Format | Size (bytes) | Quality | OQ:Z | Read-Names |
|---|---|---|---|---|
| BAM | 13,017,816,377 | N/A | N/A | N/A |
| CRAM 3.0 | 7,412,319,925 | 4,106,563,351 | 2,039,939,253 | 506,820,881 |
| CRAM 3.0 small | 7,251,254,040 | 4,099,877,724 | 2,038,910,887 | 417,349,414 |
| CRAM 3.1 | 7,110,617,135 | 4,108,014,656 | 1,954,404,510 | 295,405,271 |
| CRAM 3.1 small | 6,740,573,443 | 3,834,028,984 | 1,944,155,206 | 288,739,462 |

The original data has been passed through GATK's quality score recalibration tool, with the original qualities copied to the OQ:Z BAM auxiliary tag and the updated qualities put into the QUAL field. It can be seen that this has dramatically increased the entropy of the quality values. Note we believe this approach is no longer part of the GATK best practice, but have no direct experience in this. However the data files are a real dataset.

The default CRAM 3.0 file is 43% smaller than the default BAM, and the default CRAM 3.1 is 4.1% smaller than the 3.0.

Table 15: SynDip compression, post-Crumble

| Format | Size (bytes) | Quality | Read-Names |
|---|---|---|---|
| BAM | 3,847,978,151 | 447,519,295 | 362,906,006 |
| CRAM 3.0 | 944,003,579 | 234,945,688 | 49,496,197 |
| CRAM 3.0 small | 892,917,164 | 229,930,974 | 36,065,903 |
| CRAM 3.0 archive9 | 849,921,368 | 227,566,075 | 32,465,280 |
| CRAM 3.1 | 900,645,212 | 214,840,082 | 32,539,728 |
| CRAM 3.1 small | 862,661,875 | 212,046,488 | 27,979,249 |
| CRAM 3.1 archive9 | 817,935,498 | 203,689,827 | 23,413,440 |
| CRAM 4.0 | 891,445,170 | 215,634,168 | 29,256,258 |
| CRAM 4.0 small | 851,998,233 | 212,630,819 | 24,391,963 |
| CRAM 4.0 archive9 | 801,384,897 | 193,674,218 | 19,812,538 |

Note for the default BAM mode we estimated the space taken by quality values and read-names by replacing these fields with "*" and measuring the impact on the BAM file size.

The Crumble output here is modifying quality scores as outlined above, but also discarding read names (via HTSlibs "lossy-names" option) where the read-pair falls entirely within a single CRAM slice, and removing the large OQ:Z auxiliary tag. The modified qualities have reduced in size by 19-fold.

It can be seen that all versions of CRAM now significantly outperform BAM by a large ratio ratio than pre-Crumble, saving between 75 and 79% of storage. CRAM 3.1 is between 3.4 and 4.6% smaller than the analogous CRAM 3.0 file, mirroring the relative gains we saw in the pre-Crumble data. Also included is the early prototype for CRAM 4.0, which slightly pushes this trend further.

For both pre- and post-crumble data CRAM 3.1's default compression level is on within 1% of or smaller than CRAM 3.0's small profile (using larger block sizes and bzip2), while also being faster (speed not shown above).

### 6 Random access

High data compression is useful, but it sometimes comes with a random access cost. As we make the blocks larger than the granularity of random access coarser, the compression ratio will improve.

The comparisons so far have been with mostly default options and different tools have very different default block sizes. For The NA12878 NovaSeq data set we have computed the average number of alignment record for block as reported by the tools. These can be seen in Table 16.

Table 16: Record numbers per file block

| Format | No. Records |
|---|---:|
| BAM | 156 |
| CRAM (normal) | 10,000 |
| CRAM small | 25,000 |
| CRAM archive9 | 100,000 |
| Deez | 216,015 |
| Genozip | 39,883 |
| MPEG-G | 125,763 |

BAM record count was computed by dividing the uncompressed file by $0xff00$ (the size of a BGZF data block), and then dividing the total number of records by that figure. Similarly Deez and Genozip numbers are computed from the total record count and number of blocks. MPEG-G is reported to use 65,536 record pairs per Access Unit, and we counted the number of records observed per 65,536 uniq read identifiers. This is not quite a 2:1 ratio as some templates span access units or are aligned far away.

To test the I/O performance with random access, we performed a single range query on Chr1 at position 10,000,000 of length 1Kbp, 10Kbp, 100Kbp, 1Mbp and 10Mbp (ie position 10M to 20M). The average depth across the largest region was 36x. We used the `io_trace` tool to evaluate the number of read and seek system calls, and the total count of bytes transferred. We repeated the test using Unix `time` to measure the elapsed time to query and report SAM records, without threading enabled. These were piped into `wc -l` to validate the result. These results are presented in Table 17.

There were some disparities in the number of records reported, with Deez and Genozip reporting records that start on or after Chr1:10000000 rather than records that overlap. There is still some small discrepancy between these tools, but this was not investigated.

I/O stats are not visible for Deez as it uses memory-mapped I/O which cannot be seen in a system call trace. All reference based tools exclude the I/O on the reference itself (which is also mmaped for the HTSlib CRAM implementation). CRAM and BAM I/O figures did not include the index, while Genozip and Deez have an internal index. The index file size for BAM was 9,282,200 bytes and for CRAM it was 1,379,148 bytes.

A more complete comparison was made with BAM, CRAM 3.0 and CRAM 3.1 accessing all exon and all gene regions across Chromosome 1. The BED file was produced from Ensembl release 104 GTF at http://ftp.ensembl.org/pub/current_gtf/homo_sapiens/.

This was converted to exon and gene bed file with awk, and tested using the SAMtools multi-region iterator:

```
zcat Homo_sapiens.GRCh38.104.gtf.gz | awk '$1 == 1 && $3 == "gene" \
    {printf("chr%d\t%d\t%d\n", $1, $4-1, $5)}' > genes.bed
zcat Homo_sapiens.GRCh38.104.gtf.gz | awk '$1 == 1 && $3 == "exon" \
    {printf("chr%d\t%d\t%d\n", $1, $4-1, $5)}' > exons.bed

samtools view -f 0xffff -M -L exons.bed NA12878.final.bam
```

The genes and exons files have 5475 and 136680 regions respectively, covering 78.6Mbp and

Table 17: I/O figures per random access unit

| Format | Region | Time(s) | Records | MBytes | Reads | Seeks |
|---|---|---|---|---|---|---|
| BAM | 1Kbp | 0.066 | 284 | 0.29 | 10 | 3 |
| BAM | 10Kbp | 0.069 | 2,597 | 0.43 | 14 | 3 |
| BAM | 100Kbp | 0.115 | 26,222 | 1.71 | 53 | 3 |
| BAM | 1Mbp | 0.573 | 266,894 | 14.39 | 440 | 3 |
| BAM | 10Mbp | 6.290 | 2,693,407 | 143.23 | 4,373 | 3 |
| CRAM 3.1 | 1Kbp | 0.047 | 284 | 0.36 | 10 | 1 |
| CRAM 3.1 | 10Kbp | 0.047 | 2,597 | 0.36 | 10 | 1 |
| CRAM 3.1 | 100Kbp | 0.116 | 26,222 | 0.72 | 20 | 1 |
| CRAM 3.1 | 1Mbp | 0.823 | 266,894 | 4.95 | 145 | 1 |
| CRAM 3.1 | 10Mbp | 8.115 | 2,693,407 | 49.22 | 1,361 | 1 |
| Genozip | 1Kbp | 13.059 | 250 | 60.67 | 382 | 272 |
| Genozip | 10Kbp | 13.595 | 2,563 | 61.48 | 426 | 335 |
| Genozip | 100Kb | 13.636 | 26,188 | 61.48 | 426 | 335 |
| Genozip | 1Mbp | 17.276 | 2,66,860 | 66.42 | 708 | 712 |
| Genozip | 10Mbp | 53.834 | 26,93,373 | 114.60 | 3,476 | 4,443 |
| Deez -q1 | 1Kbp | 6.650 | 250 | N/A | N/A | N/A |
| Deez -q1 | 10Kbp | 6.537 | 2,562 | N/A | N/A | N/A |
| Deez -q1 | 100Kbp | 6.493 | 26,188 | N/A | N/A | N/A |
| Deez -q1 | 1Mbp | 7.682 | 266,859 | N/A | N/A | N/A |
| Deez -q1 | 10Mbp | 27.587 | 2,693,372 | N/A | N/A | N/A |

10.2Mbp of chromosome 1. Total I/O figures were achieved with io_trace:

```
io_trace -r ‘pwd‘ -- samtools view -f 0xffff -M -L exons.bed \
    ‘pwd‘/NA12878.final.31.cram
```

Table 18 shows the I/O statistics for the exons and genes regions.

Table 18: I/O statistics when querying many regions

| Format | Regions | Time(s) | Records | Data(MB) | Reads | Seeks |
|---|---|---|---|---|---|---|
| BAM | Exons | 30.4 | 4,894,064 | 1,489.7 | 45,503 | 5,587 |
| CRAM 3.1 | Exons | 65.6 | 4,894,064 | 844.6 | 23,017 | 580 |
| CRAM 3.1 small | Exons | 89.0 | 4,894,064 | 775.3 | 16,713 | 123 |
| BAM | Genes | 19.0 | 37,263,364 | 2,234.2 | 68,244 | 1,646 |
| CRAM 3.1 | Genes | 71.6 | 37,263,364 | 910.4 | 24,559 | 331 |
| CRAM 3.1 small | Genes | 92.7 | 37,263,364 | 792.0 | 16,991 | 88 |

The I/O load is significantly lower for the CRAM files, but elapsed time was larger. Note these test files were in disk cache, so this mainly reflects CPU time. It is unknown what the elapsed time would be after purging from cache as we were unable to perform this experiment, but queries the genes.bed file on a different BAM out of disk cache showed 85s uncached and an I/O read throughput of 45MB/s. This would likely indicate on uncached data BAM is still faster than CRAM with local storage. For remote network based I/O, for example via an htsget server, we would expect this to be reversed due to fewer access regions (seeks), higher latencies and probable lower transfer speeds..
